# MAETi: Mild acid elution in a tip enables immunopeptidome profiling from 1E4 cells

**DOI:** 10.1101/2024.12.20.628848

**Authors:** Julian Beyrle, Yuzuru Yamasaki, Megan Ford, Claire Conche, Yannic Chen, Annette Arnold, Riccardo Pecori, Ute Distler, David Gomez-Zepeda, Stefan Tenzer

## Abstract

Mass spectrometry (MS)-based immunopeptidomics is essential for characterizing MHC class I (MHC-I) immunopeptides and defining targets for tumor immunotherapies. Conventional workflows require large sample amounts, limiting their use in low-input scenarios, high-throughput screens, and kinetic studies. We developed Mild Acid Elution in a Tip (MAETi), an antibody-free, low-input method for MHC-I immunopeptidome profiling. An optimized β-alanine MAE buffer reduces background, enhances peptide coverage, and improves reproducibility. Compared with bulk immunoprecipitation, β-alanine MAE achieves similar or complementary peptide depth. Using two protocol formats we profiled 1E4-1E6 JY cells, detecting on average over 3,000 and 6,000 predicted binders, respectively. A personalized database with single amino acid variants (SAVs) enabled robust identification and quantitation of SAV epitopes from only 1E4 JY cells. Additionally, MAETi supported differential ligandome analysis of 1.5E4-5E4 FACS-sorted immune cells, revealing cell type-specific ligandome repertoires. MAETi provides a simple, fast, and scalable workflow for MS-based immunopeptidomics from minimal sample input.

## 1. Introduction

Mass spectrometry (MS)-based immunopeptidomics is a key method for understanding immune surveillance and developing immunotherapies.^1–3^ The field advances quickly and rapid technological, instrumental, and bioinformatic developments in the past 10 years have enabled astonishing progress in the field^4^. Despite these advancements, the need for high sample amounts for most workflows substantially limits translation into the clinic, where patient material for characterization and analysis is typically scarce^5,6^. Moreover, composition and abundance of the immunopeptidome change dynamically under treatment, disease, infection, or cellular stress, even within the same cell type^7–9^. Thus, unravelling immunopeptidome heterogeneities in low-input scenarios, such as sorted-cell populations, is critical for better understanding antigen presentation dynamics, immune recognition, and for designing epitope-specific immunotherapies. ^10–12^ However, this requires fast, scalable, and sensitive immunopeptidomics workflows, which until recently has been a challenge for established sample preparation methodologies.

There are two major immunopeptide enrichment strategies (Fig. 1a). Immunoprecipitation (IP) is the most commonly used approach and employs a bead-immobilized antibody to enrich MHC-peptide complexes (class I or II) from cell or tissue lysates and body fluids^4,13,14^. In contrast to antibody-based workflows, mild acid elution (MAE) takes advantage of the dissociation of MHC class I (MHC-1) complexes in mild-acidic conditions, thereby eluting MHC-1 bound peptides (MHC1ps) directly from intact cells^15,16^. In both protocols, potential MHC1ps, which are typically in the length range of 8 to 13 amino acids (8-13 mers), are further purified and analyzed by Liquid-Chromatography Mass Spectrometry (LC-MS)^16^. Historically, MAE was the first method used to isolate immunopeptides, but it has been widely replaced by IP to achieve higher enrichment specificity. More recent work by Sturm *et al.*^16^ challenged this view and demonstrated that an optimized MAE workflow can achieve immunopeptide coverage and enrichment specificity comparable to IP. These findings indicate that MAE is a viable alternative to IP. Nevertheless, both methods involve labor-intensive sample cleanup steps such as centrifugation, ultrafiltration, and offline solid-phase extraction (SPE, Fig. 1b, top) ^4,13,16^. This is a lengthy process associated with sample loss and a lowered ligandome yield, requiring high amounts of sample material^17–21^. More recently, microfluidics-based approaches have gained traction in immunopeptidomics for their ability to handle smaller sample volumes, making them well-suited for low-input samples^17,22,23^ (Fig. 1a, top). However, the requirement for specialized equipment and expertise to manufacture and operate the microfluidic chips has so far hindered broader adoption and throughput^17,22,23^.

**Fig. 1:**
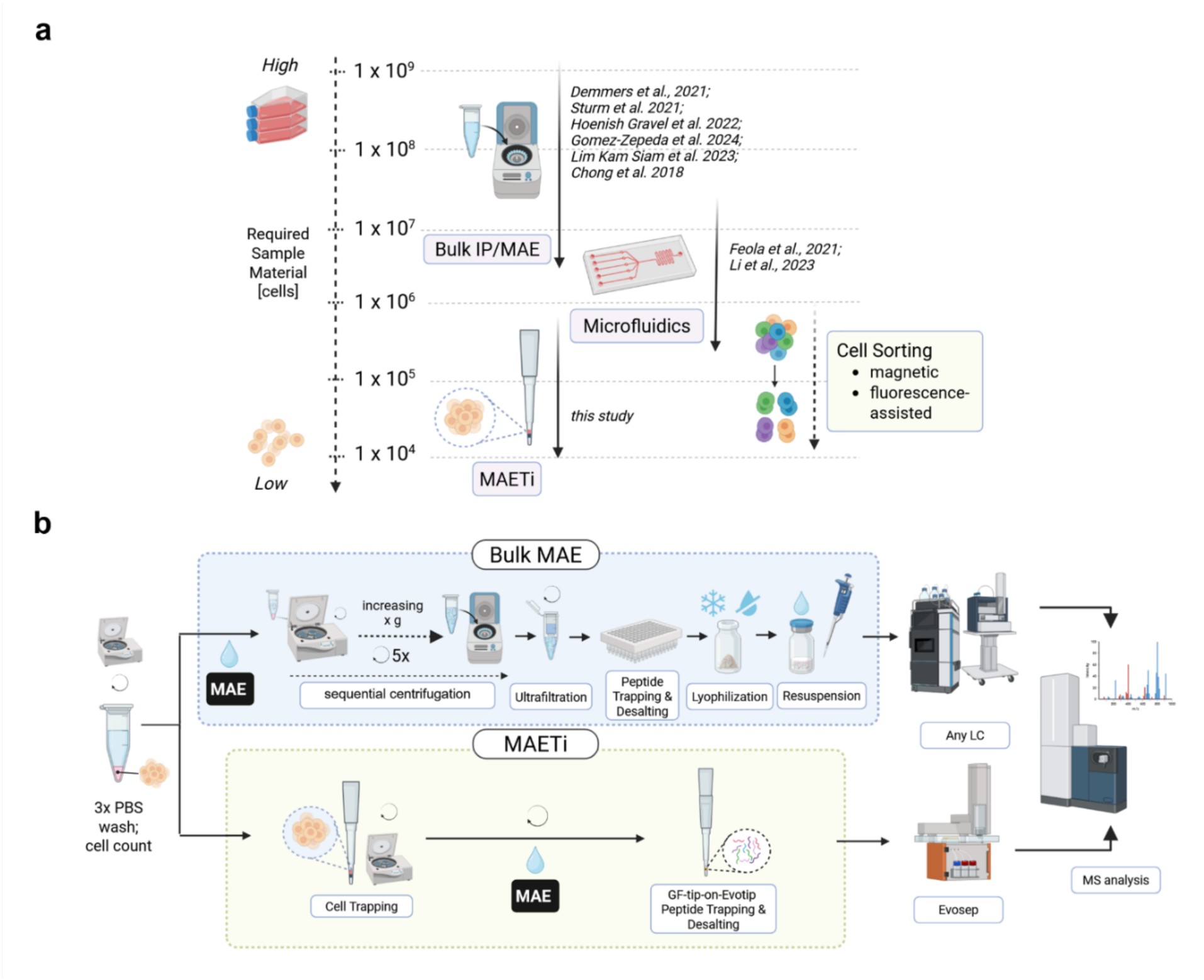
Schematic overview of current immunopeptidomics methodologies and a summary of experimental steps in Mild Acid in a Tip (MAETi) **a**, Schematic Overview showing current immunopeptidomics methodologies in the context of required sample input material, including bulk, microfluidics, and our Mild Acid Elution in a Tip (MAETi) workflow. **b**, MAETi profits from the substantial reduction of sample handling steps compared to a bulk MAE workflow and enables immunopeptidome profiling down to 1E4 cells. MAETi utilizes a glass fiber membrane tapered into a pipette tip retaining MAE-stripped cells while directly recovering eluted peptides on disposable trap columns (Evotips) prior to desalting and MS analysis. IP, immunoprecipitation; MAE, mild acid elution; FACS, fluorescence-assisted cell sorting; MACS, magnetically-assisted cell sorting. Created in BioRender. Tenzer, S. (2026) https://BioRender.com/x91a854

To provide a simple and viable alternative to complex microfluidics layouts, we have developed an optimized Mild Acid Elution in a Tip (MAETi) workflow (Fig. 1b, bottom). Briefly, pre-washed cells are placed onto glass fiber (GF) membrane-tapered pipette tips and peptides are directly stripped by MAE onto disposable trap columns (Evotips). This eliminates most steps required in bulk immunopeptidomics enrichment workflows^24–26^, improves overall sensitivity, and enables direct ligandome analysis down to 1E4 cells. After desalting on the Evotip, the MAE-eluate is injected onto a reverse-phase column using the Evosep One liquid-chromatography (LC) system coupled to a timsTOF mass spectrometer.

We present MAETi, a minimal, simplified sample preparation protocol enabling robust MHC-1 ligandome analysis using 1E4 to 1E6 cells per sample as an initial input. Elution buffer optimizations resulted in cleaner samples and boosted 8-13mer identifications by over 50% compared to traditional buffer formulations. By applying a personalized database and MAETi with diaPASEF we robustly quantify single-amino acid variants (SAVs) down to 1E4 JY cells. Given the low cell input required, we show that MAETi enables differential ligandome analysis of 1.5E4-5E4 healthy-donor-derived immune cells sorted by fluorescence-activated cell sorting. This technology has the potential to resolve heterogenous immunopeptidomes from low-abundant cell populations (e.g., under the influence of the tumor microenvironment) or facilitate kinetic profiling of drug-treated cells. Thus, MAETi holds significant potential for advancing tumor immunology, personalized immunotherapy, and drug discovery.

## 2. Results

### 2.1. A minimal MAE-based MHC1p enrichment workflow enables immunopeptidomics profiling from 5E4 cells

To demonstrate the feasibility of our MAETi protocol (Fig. 1b, bottom), we processed individual aliquots of 5E4 JY cells. Captured peptides of 5E4 cells per Evotip were separated using the 40 samples per day (SPD) Whisper gradient, and mass spectrometry data were acquired on a timsTOF SCP (Fig. 2a) using ddaPASEF with two distinct isolation polygons (n=3 each) (Fig. 2b). This stepped precursor isolation polygon (Fig. 2b-left) was designed to fragment all possible peptides covering the 1/K_0_ vs. *m/z* ion clouds that correspond to all multiply and singly charged ions above *m/z* 600. In contrast, Thunder-ddaPASEF (Fig. 2b-right) efficiently uses the instrument time by focusing the MS/MS events on the ion cloud corresponding to 8-13mers^27^.

**Fig. 2:**
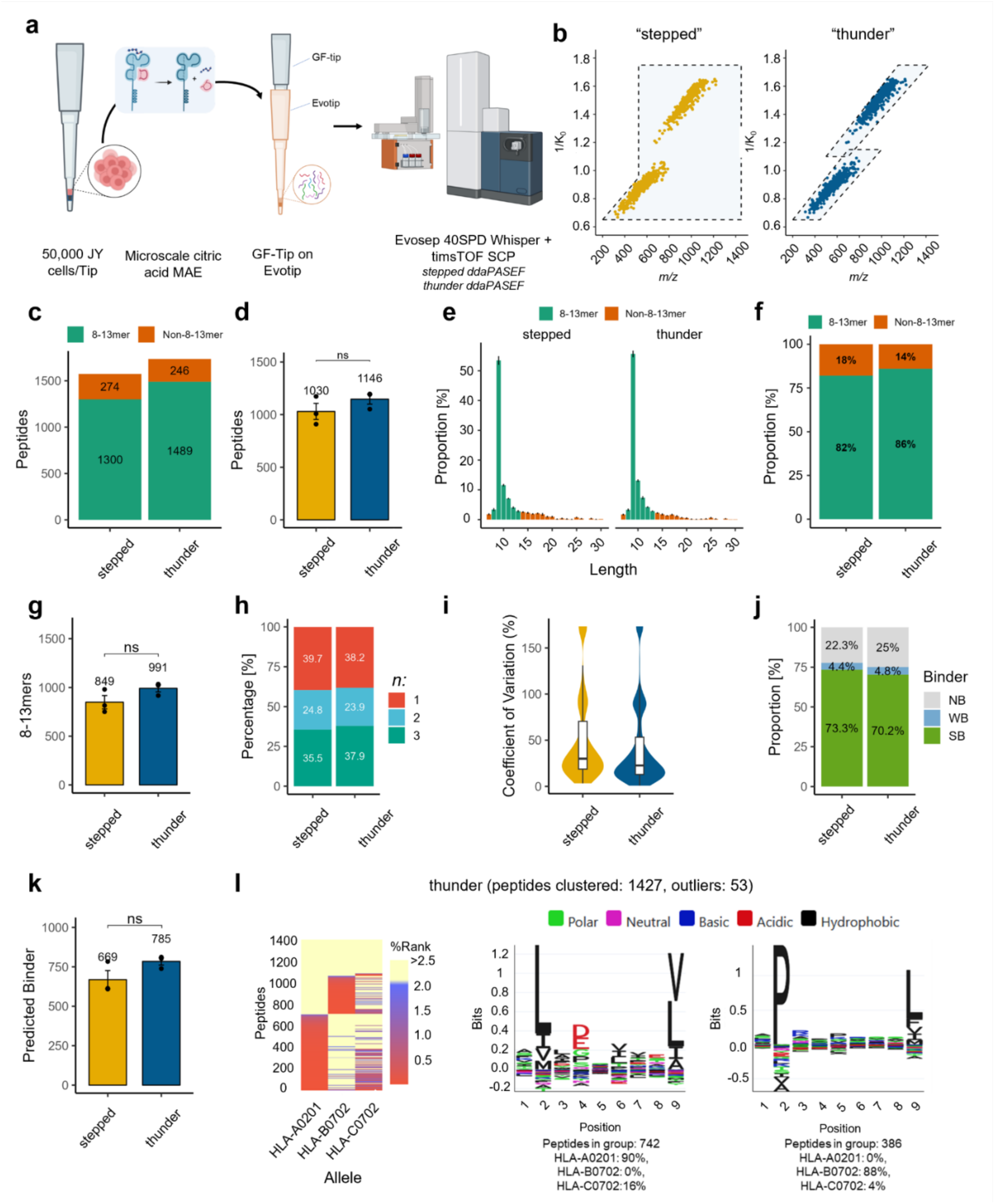
MAETi enables ligandome analysis of 5E4 JY cells. **a-b,** Workflow of Mild Acid Elution in a Tip (MAETi) proof of principle experiment and DDA acquisition schemes used. **a,** Samples of 5E4 JY cells each were prepared by MAETi, directly loaded onto Evotips, and analyzed in an Evosep One coupled to a timsTOF SCP, using a Whisper 40 SPD gradient and two distinct DDA acquisition schemes (n=3, each). *Created in BioRender. Tenzer, S. (2026)* https://BioRender.com/u18z153. **b** 1/K_0_ (inverse ion mobility, invkv0) vs. m/z peptide distribution showing the polygons used for unbiased Stepped-ddaPASEF and MHC-tailored Thunder-ddaPASEF; dots represent identified 8-13mers. **c-i** Experimental characteristics of MHC-1 immunopeptide enrichment using MAETi. **C,** Total peptides identified per acquisition scheme across three injection replicates, divided into 8-13mers and other peptide lengths. **d,** Peptides identified on average per sample (n=3). **e,** Histograms showing the peptide length distribution for both acquisition schemes. **f,** Proportion of peptides with the expected peptide length for MHC class I ligands (8-13mer). **g,** Peptides with the expected size (8-13mers) identified on average per sample. **h,** Data completeness shown as the proportion of 8-13mers identified in one, two, or all the three replicates (n). **i,** Violin and boxplots of the peak area coefficient of variation (CV) of 8-13mers identified in all 3 replicates (center line, median; box limits, upper and lower quartiles; whiskers, 1.5x interquartile range). **j,** Proportion of 8-13mers predicted to bind JY MHC alleles (NB, non-binders; WB, weak binders; SB, strong binders). **k,** MHC predicted binders (unique SB and WB) identified on average per sample. **I,** Unsupervised sequence clustering of the peptides identified, matching the expected motifs for JY MHC alleles. Bars with error bars correspond to *average ± sd* with points indicating the individual values, and asterisks designate statistical differences (two-sided t-test; * *p* = 0.05, ** *p* = 0.01, *** *p* = 0.001).

First, we assessed the peptide length distribution using an unbiased approach (Stepped-ddaPASEF). This resulted in 1574 total peptides (Fig. 2c) identified across three replicates, with 1,030 ± 133 identifications per sample (mean ± sd) (Fig. 2d). The enriched peptides exhibited the expected length distribution for MHC1ps with an apex at 9mers (Fig. 2e) and were predominantly 8-13mers (82%, Fig. 2f), corresponding to 849 ± 118 8-13mers identified on average per sample (mean ± sd; Fig. 2g). These results indicate that MAETi enables the direct identification of peptides with an MHC-1 -typical peptide length distribution and does not lead to co-enrichment of longer peptides.

Next, we implemented the MHC1p-tailored thunderbolt-shaped polygon into the Thunder-ddaPASEF method (Fig. 2b-right^27^), which resulted in a 10% increase in peptide identifications, both in total and on average (Fig. 2c, 2d) compared to the stepped polygon. In addition, the thunder polygon slightly increased the number of 9mers (Fig. 2e) and the proportion of 8-13mers (from 83% to 86%; Fig. 2f). Data completeness was comparable between the stepped (36%) and thunder polygon (38%; Fig. 2h). However, DDA acquisition using the Thunder isolation polygon increased the quantitative precision as indicated by a decreased median coefficient of variation (CV) of the precursor (MS1) area of 8-13mers from 30% to 22% (Fig. 2i).

Finally, to verify the capability of MAETi to enrich MHC1ps, we used MHCvizPipe^28^ to predict peptide binding to JY MHC alleles (HLA-A*02:01, HLA-B*07:02, and HLA-C*07:02 via NetMHCPan-4.1^29^) and generate unsupervised and supervised sequence clustering analyses (via GibbsClustering^30^). Although both MS methods yielded nearly identical proportions of predicted binders (strong and weak binders, Fig. 2j), the Thunder-ddaPASEF detected more predicted binders on average than the stepped polygon (*p* = NS; Fig. 2k). Despite the different isolation polygons, both methods shared high overlaps for peptides (58.4%), 8-13mers (59.4%) and predicted binders (64.7%) demonstrating unbiased MHC1p enrichment (Supplementary Fig. S1a-c). Importantly, in both cases, unsupervised Gibbs Clustering of the enriched peptides displayed the expected amino acid sequence motifs previously described in other studies for the JY MHC-Alleles, including IP-based enrichments^16,27^ (Fig. 2l for Thunder-ddaPASEF and Supplementary Fig. S1d for both MS-methods; allele-supervised Gibbs Clustering for both MS-methods Supplementary Fig. S1e).

These results demonstrate that MAE of whole cells trapped on a glass-filter tip can be used to enrich the MHC class 1 immunopeptidome. This initial version of MAETi enabled the MHC1p immunopeptidomics profiling of as few as 5E4 cells, yielding on average 785 ± 40 predicted binders on a timsTOF SCP instrument using Thunder-ddaPASEF.

### 2.2. Optimized β-alanine-based MAE buffer provides cleaner samples and enhances immunopeptidome coverage and reproducibility

High-sensitivity immunopeptidome profiling of minute sample amounts requires high-purity reagents to minimize the risk of introducing contaminants during sample preparation. After initial validation of the citric acid-based MAETi protocol, closer inspection of the raw data revealed a high background level in the acquired LC-MS/MS datasets. To reveal the source of this potential contamination, we analyzed pure (≥99%) 131 mM citric acid (pKa ∼ 3.13) loaded into Evotips (Fig. 3a-top). The resulting data showed up to 7-fold higher base peak chromatogram (BPC) intensities than a blank injection of 0.1% formic acid (FA) (Fig. 3b), with several highly intense ions eluting across the whole gradient (Fig. 3c). These results clearly indicate, that citric acid introduced a high level of background ions, which potentially interfere with analyte ionization despite extensive washing of the Evotips (3x 0.1% FA in water).

**Fig. 3:**
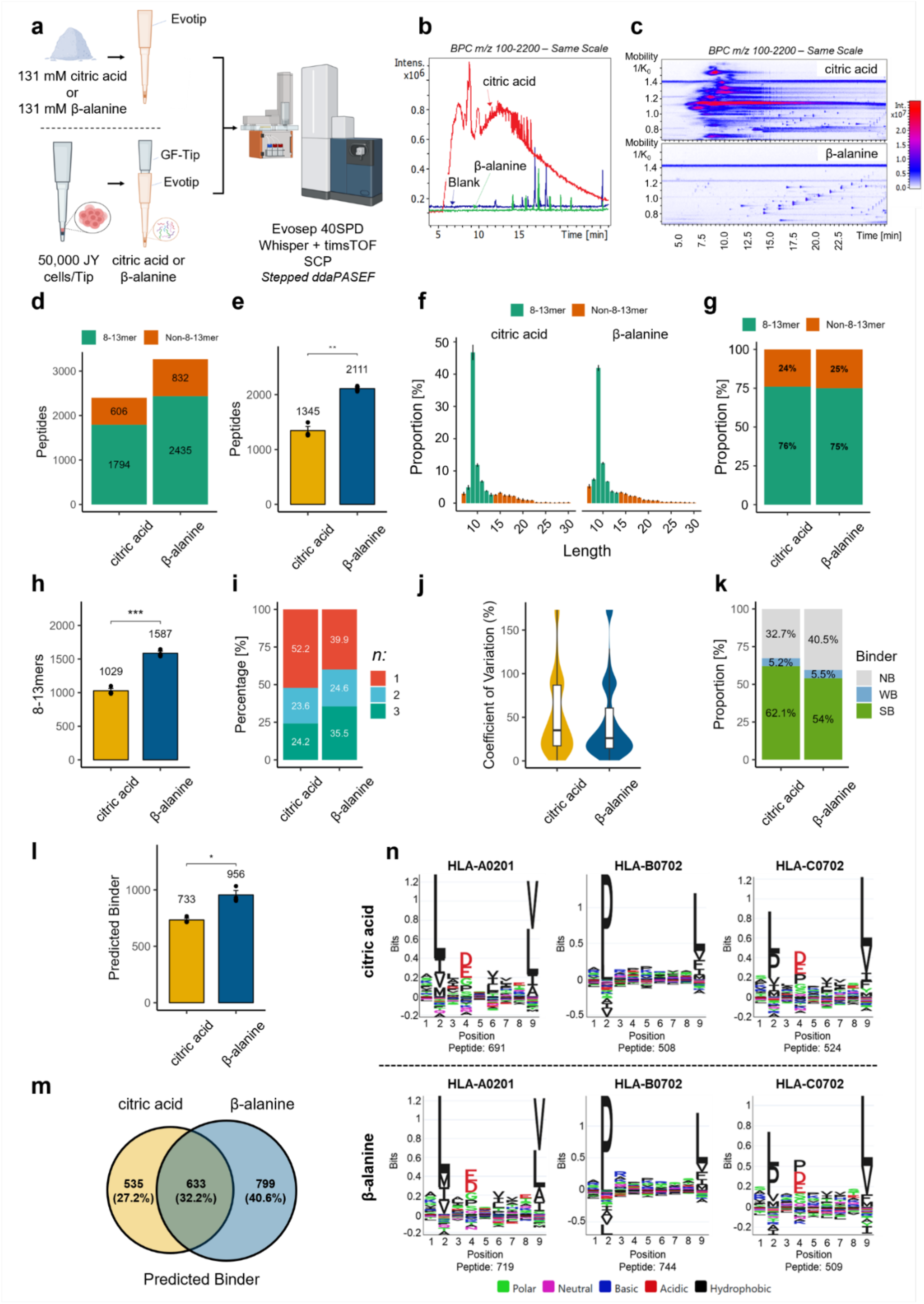
Alternative MAETi buffer composition results in cleaner MS spectra and improves ligandome recovery. **a-n** Experimental layout for MAE buffer optimizations (**a**), testing pure buffer substances (**b-c**) and benchmarking alternative buffer performance for MAETi analysis of JY cells (**d-n**). **a** Either 131 mM citric acid or β-alanine were directly loaded into Evotips to assess background contaminant levels; Then whole MAETis (n=3) using either a citric acid- or β-alanine-based MAE buffer and with 5E4 JY cells per tip were analyzed in an Evosep coupled with timsTOF SCP using Stepped-ddaPASEF. *Created in BioRender. Tenzer, S. (2026)* https://BioRender.com/f39o469. **b** Base peak chromatograms (BPC) of 131 mM citric acid, β-alanine, or a blank (0.1% FA in water) directly loaded into Evotips. **c** Corresponding heatmaps of the MS1 ions projected in the inversed IMS vs. m/z space. **d** Total peptides identified per buffer substance across three replicates of JY cells, divided in 8-13mers and other peptide lengths. **e** Peptides identified on average per sample. **f** Histograms showing peptide length distributions for both buffer substances. **g** Proportion of peptides with the expected peptide length for MHC class I ligands (8-13mer). **h** Peptides with the expected size (8-13mers) identified on average per sample. **i** Data completeness shown as the proportion of 8-13mers identified in one, two, or all the three replicates (n). **j** Quantitative accuracy shown as violin and boxplots of the peak area coefficient of variation (CV) of 8-13mers identified in all 3 replicates (center line, median; box limits, upper and lower quartiles; whiskers, 1.5x interquartile range). **k-n** Binding predictions and sequence motifs. **k** Proportion of 8-13mers predicted to bind JY MHC alleles (NB, non-binders; WB, weak-binders; SB, strong binders). **l** MHC predicted binders (unique SB and WB) identified on average per sample. **m** Venn diagram showing the overlap of predicted binders identified in DIA (n=3) for citric acid and β-alanine. **n** Supervised Gibbs sequence clustering of the peptides identified, matching the expected motifs for JY MHC alleles. Bars with error bars correspond to *average ± sd* with points indicating the individual values, and asterisks designate statistical differences (two-sided t-test; * p ≤ 0.05, ** p ≤ 0.01, *** p ≤ 0.001, **** p ≤ 0.0001).

Therefore, we explored β-alanine as an alternative substance buffering at the pH used for MAE, with a pKa of ∼ 3.55. Notably, the analysis of pure (≥99%) β-alanine resulted in cleaner chromatographic traces with an average BPC around 8-fold lower than for citric acid, showing only a few distinct peaks (Fig. 3b). Additionally, the 1/K_0_ versus retention time (RT) heatmaps of pure β-alanine were substantially cleaner than the heatmaps of pure citric acid (Fig. 3c).

Subsequently, we assessed the suitability of β-alanine as an alternative buffer substance for MHC1p enrichment in the MAETi protocol. To this end, we prepared MAETi samples with 5E4 JY cells each using MAE buffers containing either citric acid or β-alanine at the same concentration (131 mM, n=3 per buffer) and analyzed them using the 40SPD Whisper gradient and the stepped ddaPASEF method on a timsTOF SCP mass spectrometer (Fig. 3a-bottom).

In comparison to citric acid, β-alanine yielded 36% more peptides in total and 57% more peptides on average (Fig. 3d, e, *p* = 0.01). Both buffers resulted in the expected MHC1p length distribution with > 40% 9mers (Fig. 3f) and > 75% 8-13mers (Fig. 3g). Notably, β-alanine yielded 54% more 8-13mers than citric acid (1587 vs. 1029; Fig. 3h; *p* = 0.001). In addition, β-alanine improved data completeness across sample preparation replicates (Fig. 3i), and decreased the median CV for the 8-13mer area from 35% to 26% (Fig. 3j). This indicates that peptide enrichment is more robust and reproducible with the β-alanine-based MAE, compared to citric acid.

Despite a lower proportion of 8-13mers predicted to bind JY MHC alleles (Fig. 3k) in samples prepared with the β-alanine-based MAE buffer, compared to the citric acid-based MAE buffer, around 30% more predicted binders were identified on average (956 ±68 versus 733 ±29; *p* = 0.05, Fig. 3l) for the β-alanine-based MAE buffer. Interestingly, only 32% the predicted binders were shared identifications comparing both buffers (Fig3m; Supplementary Fig. S2a-f: peptides, 8-13mers, binders). The distinct peptides for MAETi using either the citric acid or β-alanine buffer system exhibited similar peptide sequence motifs (Fig. 3n; Supplementary Fig. S2g), showing that their amino acid composition was alike. Thus, the differences could be due to biological, preparation, and analytical variations.

Altogether, these experiments demonstrate that replacing citric acid with β-alanine in the MAE buffer results in lower background ions and improves the MHC1p coverage and reproducibility of our workflow. Hence, β-alanine-based MAETi robustly identifies almost 1,000 predicted binders from only 5E4 JY cells. To further assess the quality of MHC1p enrichment using a β-alanine MAE buffer, we compare bulk β-alanine MAE buffer enrichments to bulk immunoprecipitation in the next section.

### 2.3. β-alanine-based MAE provides a complementary immunopeptidome coverage compared with immunoprecipitation

Next, we assessed the performance of the β-alanine MAE buffer and its influence on MHC1p enrichment by comparing bulk β-alanine-based MAE with bulk immunoprecipitation (IP) using the anti-pan-HLA class 1 antibody W6/32 (Fig. 4a). We profiled the two lymphoblastoid human B-cell lines JY and Raji covering 6 distinct MHC alleles and low immunopeptidome overlap (<2%, Gomez-Zepeda *et al.*^27^). Briefly, three preparation replicates of JY and Raji cells (at 4×10^7^ cells each) were prepared using MAE and IP in parallel, and each sample was injected in triplicate (2×10^6^ cell equivalents/injection, n=9 total injections). For this experiment, all samples were ultrafiltered, desalted, and analyzed on a nanoElute 2 LC system coupled to a timsTOF Pro 2 mass spectrometer using stepped ddaPASEF with settings previously optimized for this instrument (Gomez-Zepeda *et al.*^27^).

**Fig. 4:**
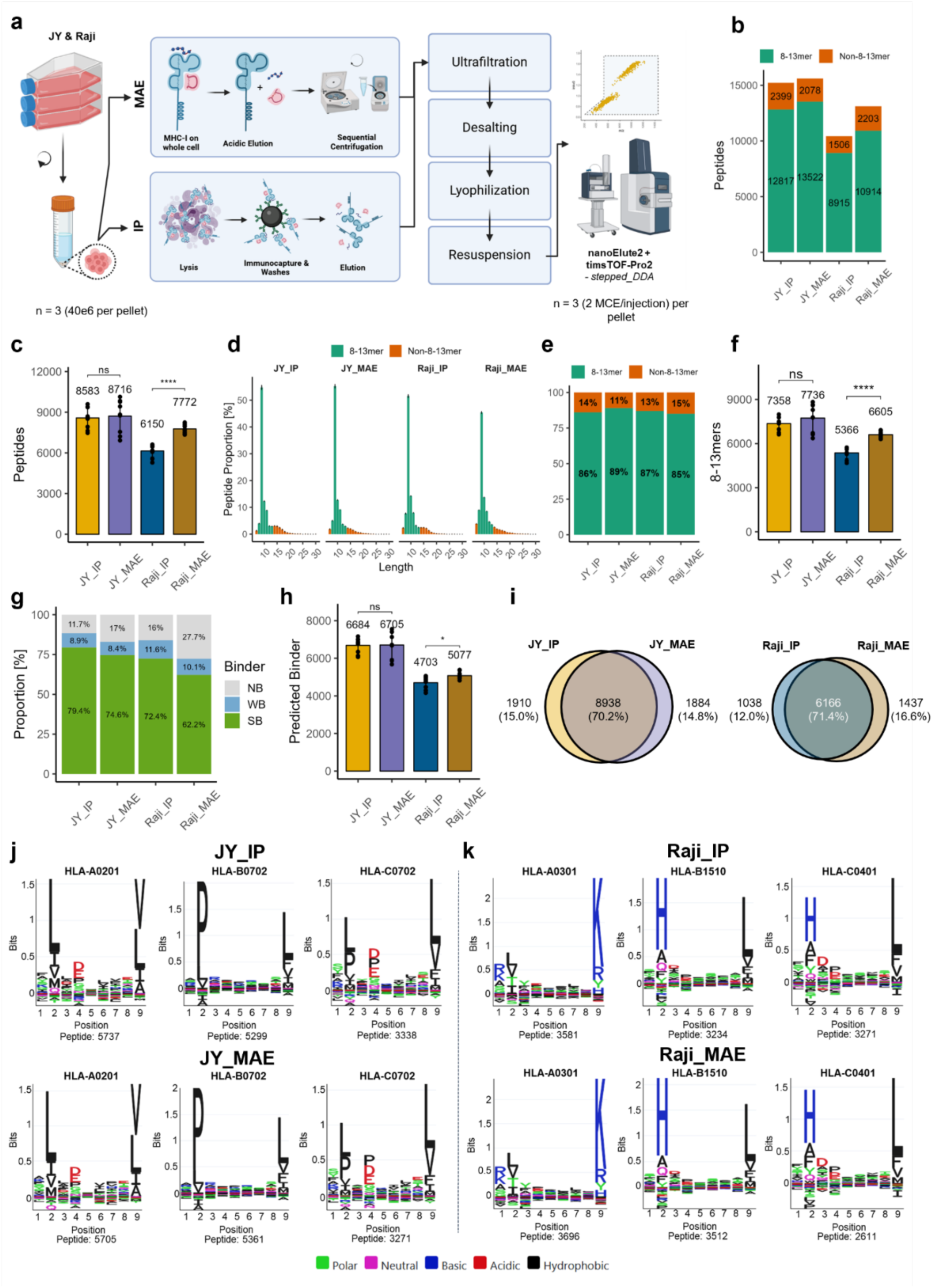
β-alanine-based MAE provides a complementary immunopeptidome coverage compared with IP in bulk experiments. **a** Experimental layout for bulk MHC-1 ligand enrichment from JY and Raji cells using MAE or IP, analyzed on a nanoElute 2 LC system coupled to a timsTOF Pro 2 mass spectrometer using Stepped-ddaPASEF. *Created in BioRender. Tenzer, S. (2026)* https://BioRender.com/i31f347. **b** Total number of peptides identified per enrichment method and cell line across three preparation and injection replicates (n=9), divided in 8-13mers and other peptide lengths. **c** Peptides identified on average per sample. **d** Histograms showing the peptide length distribution. **e** Proportion of peptides with the expected peptide length for MHC-1 ligands (8-13mer). **f** Peptides with the expected size (8-13mers) identified on average per sample. **g** Proportion of 8-13mers predicted to bind JY MHC alleles (NB, non-binders; WB, weak-binders; SB, strong binders). **h** Predicted binders identified on average per sample. **i** Venn diagram showing the overlap of binders identified in MAE and IP for JY (left) and Raji (right). **j** Supervised Gibbs sequence clustering of the peptides identified in both protocols, matching the expected motifs for the corresponding MHC alleles JY (left), and Raji (right). Bars with error bars correspond to *average ± sd* with points indicating the individual values, and asterisks designate statistical differences (two-sided t-test; * p ≤ 0.05, ** p ≤ 0.01, *** p ≤ 0.001, **** p ≤ 0.0001).

For JY cells, bulk MAE and IP resulted in similar numbers of peptides both in total (15,600 and 15,216, respectively) and on average per sample (8,716 ± 1,210 and 8,583 ± 783, respectively). However, in the case of Raji cells, MAE outperformed IP by yielding 26% more peptides in total (13,117 and 10,421, respectively) and on average (7,772 ± 324 and 6,150 ± 456, respectively, *p* ≤ 0.0001) (Fig. 4b, c). The peptide length distribution remained comparable across both methods for JY and Raji (85% to 89% 8-13mers) (Fig. 4d, e). Thus, MAE produced a significantly higher 8-13mer yield for Raji (6,605 ± 213 versus 5,366 ± 383, *p* ≤ 0.0001), but not for JY (Fig. 4f, Supplementary Fig. S3a). Incidentally, the median MS1 Area CV, dynamic range, and data completeness for 8-13mer detection across four replicates were comparable between IP and MAE within each cell line (Supplementary Fig. S3b-d).

Although the proportion of predicted binders among all identified 8-13mers was slightly lower for MAE than in IP for both JY and Raji (Fig. 4g), the average number of identified predicted binders was similar for JY and slightly higher for Raji using MAE as compared to IP (Fig. 4h, *p* ≤ 0.05). Furthermore, the overlap of predicted binders identified by MAE and IP was over 70% for both JY and Raji (Fig. 4I), with similar peptide sequence motifs obtained by supervised Gibbs Clustering (Fig. 4j-k). The similarities and differences in the peptide sequences - such as motifs, isoelectric points, or hydrophobicity - are provided in the Supplementary Fig. S3e-k.

Despite the distinct alleles and peptide motifs for Raji and JY, we observed a slight overlap in 8-13mers (∼2.5%, Supplementary Fig. S3h) but not in predicted binders (Supplementary Fig. S3i), suggesting minimal shared non-binder peptides eluted by MAE. However, this did not negatively impact predicted binder identification (Fig. 4h). Comparison of these sequence motifs (9mers, without clustering) of shared or unique identifications for MAE or IP showed no apparent motif differences for both cell lines (Supplementary Fig. S3j-k).

Collectively, these findings indicate that our β-alanine MAE results in a similar and complementary enrichment of MHC class I ligands compared to IP, rendering it a cost-effective alternative to antibody-based enrichment methods.

### 2.4. MAETi enables quantitative MHC1p-ligandome profiling of single-amino acid variants (SAVs) as a proxy for neoepitopes from 1E4 to 1E6 cells

To evaluate the quantitative performance of the optimized MAETi protocol across different cell number inputs and sample preparation layouts (GF-Tip or 96-well plate), we processed samples ranging from 10,000 (1E4) to 1,000,000 (1E6) JY cells (Fig. 5a). Samples from 1E4 to 5E4 input cells were MAE-stripped directly on the GF-Tip (on-tip), while samples from 1E5 to 1E6 input cells were stripped on a 96-well plate (on-plate).

**Fig. 5:**
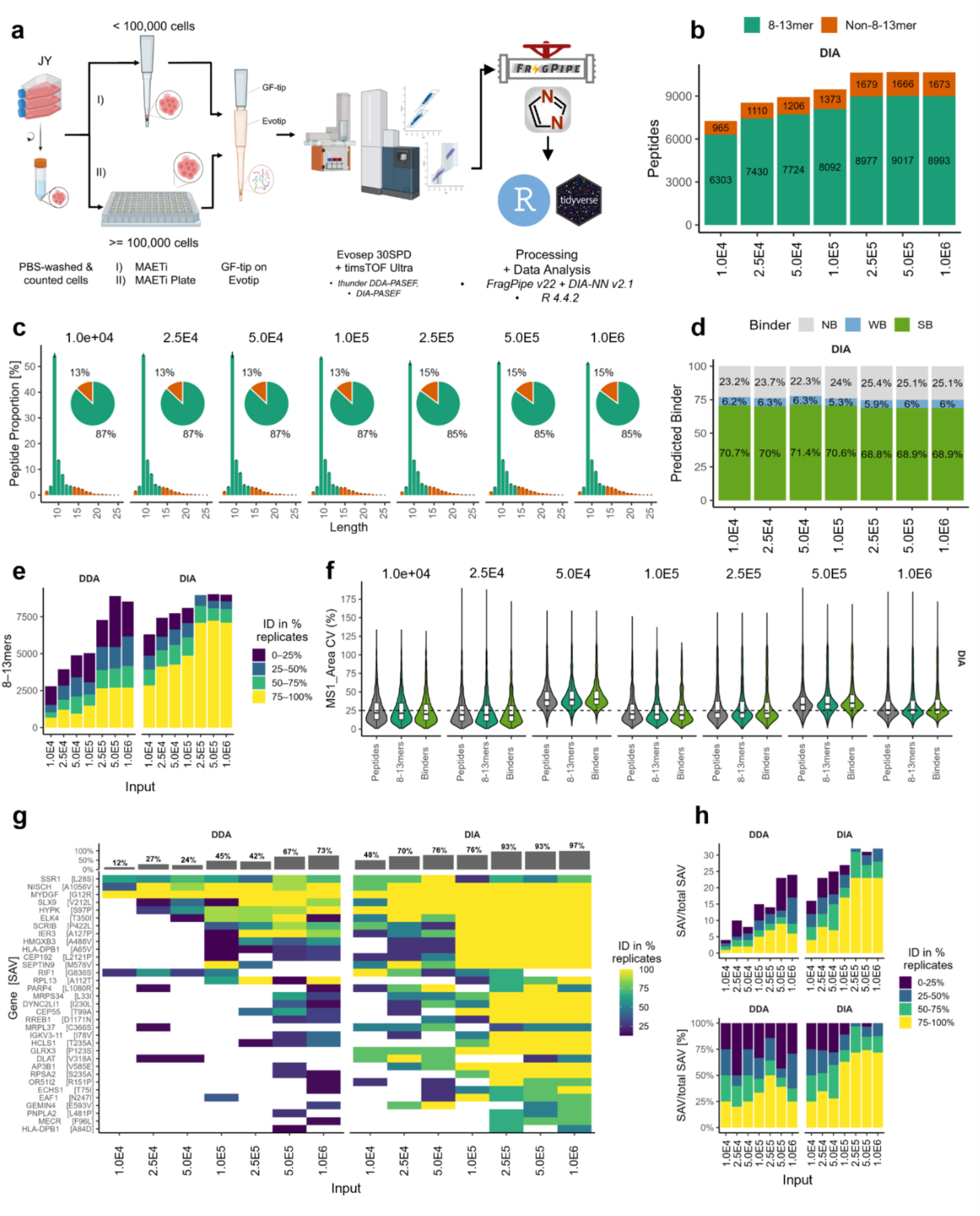
MAETi enables immunopeptidome profiling of single amino acid variants (SAVs) from 1E4 (10k) to 1E6 (1M) JY cells per sample. **a** MHC-1 ligands enriched from different input cell amounts of the human lymphoblastoid cell line JY. For cell input amounts < 1E5 (100k), cells were MAE-stripped directly on the GF-Tip onto the Evotip (**I** - on-tip format); For cell input amounts utilizing 1E5 (100k) to 1E6 (1M) cells, cells were MAE-stripped on a 96-well plate, and after centrifugation, supernatants were passed through the GF-Tip on Evotip setup (**II** - on-plate format). Loaded and washed Evotips were analyzed using an Evosep (30SPD) LC system coupled to a timsTOF Ultra mass spectrometer in Thunder-ddaPASEF and diaPASEF. *Created in BioRender. Tenzer, S. (2026)* https://BioRender.com/d35v846. **b** Unique peptides identified in total per DIA sample (divided in 8-13mers and other peptide lengths). **c** Peptide length distributions with proportion of 8-13mers and non-8-13mers **d** Proportion of 8-13mers predicted to bind JY MHC alleles (NB, non-binders; WB, weak-binders; SB, strong binders). **e** Data completeness shown as the proportion of identified 8-13mer across multiple replicates of one input cell number. **f** Distribution of mean area coefficient of variation for peptides, 8-13mers and predicted binders across the varying input numbers **g** single amino acid variant (SAV) peptides detected across the different input numbers colored by identification rate (% detected across replicates of one condition) as well as bar charts (on top) featuring proportion of SAV detection per total detected SAVs in ddaPASEF and diaPASEF. **h** Data completeness stacked bar charts for total (top) and proportional (bottom) SAV identification across the different input comparing ddaPASEF and diaPASEF.

Captured peptides were separated using the 30SPD gradient on an Evosep One LC system coupled to a timsTOF Ultra mass spectrometer. To assess both coverage and quantitative performance, we acquired the data using Thunder-ddaPASEF (n=8 each) and MHC-tailored diaPASEF^31,32^ (n=4 each). Raw files were processed in FragPipev22^33–36^ using the “Nonspecific HLA-DIA” workflow, which by default has an integrated rescoring module (MSBooster). DIA runs were loaded as DIA-Quant files and searched using the integrated DIA-NN^37,38^ (version 2.1 Academia) with a spectral library generated from the DDA runs. To enable detection of SAV peptides, a custom WES-derived JY database was used. Due to the nature of a titration experiment, non-normalized precursor quantities were used for evaluation.

As expected, the number of detected peptides and 8-13mers increased in function of the sample input in both DDA and DIA. Unique identifications ranged from a total of 2,789 to 8,530 8-13mers from 8 DDA runs per input cell number and from a total of 6,303 to 8,993 8-13mers from 4 DIA runs per input cell number from 1E4 to 1E6 cells, respectively (Fig. 5b & Supplementary Fig. S4a). However, identifications reached a plateau between 5E5 and 1E6 cells for DDA and 2.5E5 to 1E6 cells in DIA (Fig. 5b, Supplementary Fig. S4a). Importantly, ≥84% of the identified peptides were 8-13mers across all cell inputs, regardless of the sample preparation layout and acquisition method used (Fig. 5c, Supplementary Fig. S4b).

Average identifications in DIA ranged from 5,169 ± 179, to 9,573 ± 184 (*mean ± sd*) peptides, 4,491 ± 122, to 8,161 ± 168 (*mean ± sd*) 8-13mers and 3,454 ± 95 (*mean ± sd*) to 6,297 ± 164 (*mean ± sd*) predicted binders (Supplementary Fig. S4c) as sample inputs increased from 1E4 to 1E6 JY cells, respectively. Although the proportion of predicted binders (strong binders and weak binders) was slightly lower compared to immunoaffinity purified samples (see Fig. 4g), the range remained high between 67.7% to 80.5% in DDA and 74.9% to 77.7% in DIA (Fig. 5d, Supplementary Fig. S4e).

DIA provided deeper coverage than DDA, particularly at lower input levels, and good quantitative performance for all peptides, 8-13mers, and predicted binders. The average identifications were ∼3-fold higher in DIA than DDA for ≤ 2.5E4 cells, and ∼2-fold higher from 2.5E5 to 1E6 cells (Supplementary Fig. S4d). In addition, DIA improved data completeness, with up to 75% of 8-13mers reproducibly identified in 75-100% of the replicates, compared with ≤ 35% in DDA (Fig. 5e). The same applied for predicted binders (Supplementary Fig. S4f). Moreover, the median area CV across replicates was below 50% at all input levels, and below 25% in most cases (Fig. 5f). Altogether, this shows that MAETi with DIA provides deep coverage and good quantification performance down to input amounts of 1E4 cells.

Next, we complemented the canonical database (Uniprot human proteome FASTA download 2023-6-29) using variant calling information from WES data to identify potential SAVs as a proxy for neoepitopes and to assess the quantitation in function of decreasing input cell numbers. Proteins featuring >2 SAVs/protein and thus likely originating from highly polymorphic genes (e.g., HLA-A with 18 SAVs/protein) were excluded from the identified peptides.

Among the 11,243 uniquely identified peptides in the dataset, 33 represented SAVs. Next, we evaluated the percentage of SAVs consistently detected across all replicates within a given cell input and acquisition mode (Fig. 5g). Compared to DDA, DIA identified more SAV peptides, proving its higher sensitivity for potential neoantigen detection (Fig. 5g). More precisely, in DDA 12% (4/33) of the SAVs were detected from 1E4 JY cells, and >50% (22/33) identified SAVs were reached at 5E5 cells (Fig. 5g, bar charts, top). In contrast, DIA identified almost 50% of the SAV peptides at all the input levels, from as few as 1E4 cells, and reached 93% from 2.5E5 to 1E6 cells. This shows that MAETi in combination with diaPASEF enables high-sensitivity detection of SAV peptides or potential neoantigens from as few as 1E4 cells.

In summary, these results show that MAETi can be applied from 1E4 cells on-tip and up to 1E6 cells on-plate (96-well plate) to enrich MHC1ps with high selectivity. We further demonstrate that MAETi using diaPASEF enables robust detection of SAVs down to a minimal input number of 1E4 JY cells. Furthermore, the advantages of DIA, including its increased sensitivity, dynamic range, and improved data completeness for MHC1p identifications^39^ make the workflow particularly well-suited for robust quantitative profiling of MHC1p-ligandomes across a wide range of cell inputs. This renders MAETi a versatile tool for immunopeptidomics analyses of limited sample amounts.

### 2.5. MAETi enables MHC1 ligandomics of FACS-sorted immune cells

The immunopeptidome mirrors the cellular state, with composition varying across cell types and abundance shifting dynamically even within the same cell type in response to treatment, disease, infection, or stress.^24^ Thus, characterizing the heterogeneity of MHC-1–presented peptides across cell types and determining which potential MHC1ps increase or decrease for a given cell type and stimulus is critical for understanding antigen presentation dynamics, immune recognition, and designing immunotherapies.^10–12^

To investigate the ability of MAETi to resolve the immunopeptidome of sorted cell populations, human peripheral blood mononuclear cells (PBMCs) from four healthy donors (D1-D4) were subjected to fluorescence-assisted cell sorting (FACS, Supplementary Fig. S5a). FACS-isolated cell populations (HLA-ABC^+^ & CD14^+^, CD3^+^ and CD16^+^CD56^+^) were MAE-stripped on-tip at 1.5E4 (15k), 2E4 (20k) and 5E4 (50k) cells, acquiring up to four replicates per cell type using diaPASEF.^32^ FACS counts/events were used as a direct proxy for cell numbers. Whole purified unsorted PBMCs were MAE-stripped on-plate at 2.5E5 (250k, D1-4), 5E5 (500k, D2-4), and 7.5E5 (750k, D1) cells and measured in Thunder-ddaPASEF to create donor-specific spectral libraries. Unsorted JY cells (5E4 cells, n=2) were prepared in parallel as quality control. Samples were analyzed using an Evosep (30SPD) LC system coupled to a timsTOF Ultra mass spectrometer in Thunder-ddaPASEF and diaPASEF. Data derived from each donor was processed individually in Fragpipev23.0^33,35,36,40^ automatically generating donor-specific, DDA-based PBMC spectral libraries (n_2.5E5_ = 3-4, n_5E5_ = 4, n_7.5E5_ = 3) for cell-specific DIA (DIAQuant) searches using DIA-NNv2.1^37,38^ (Fig. 6a).

**Fig. 6:**
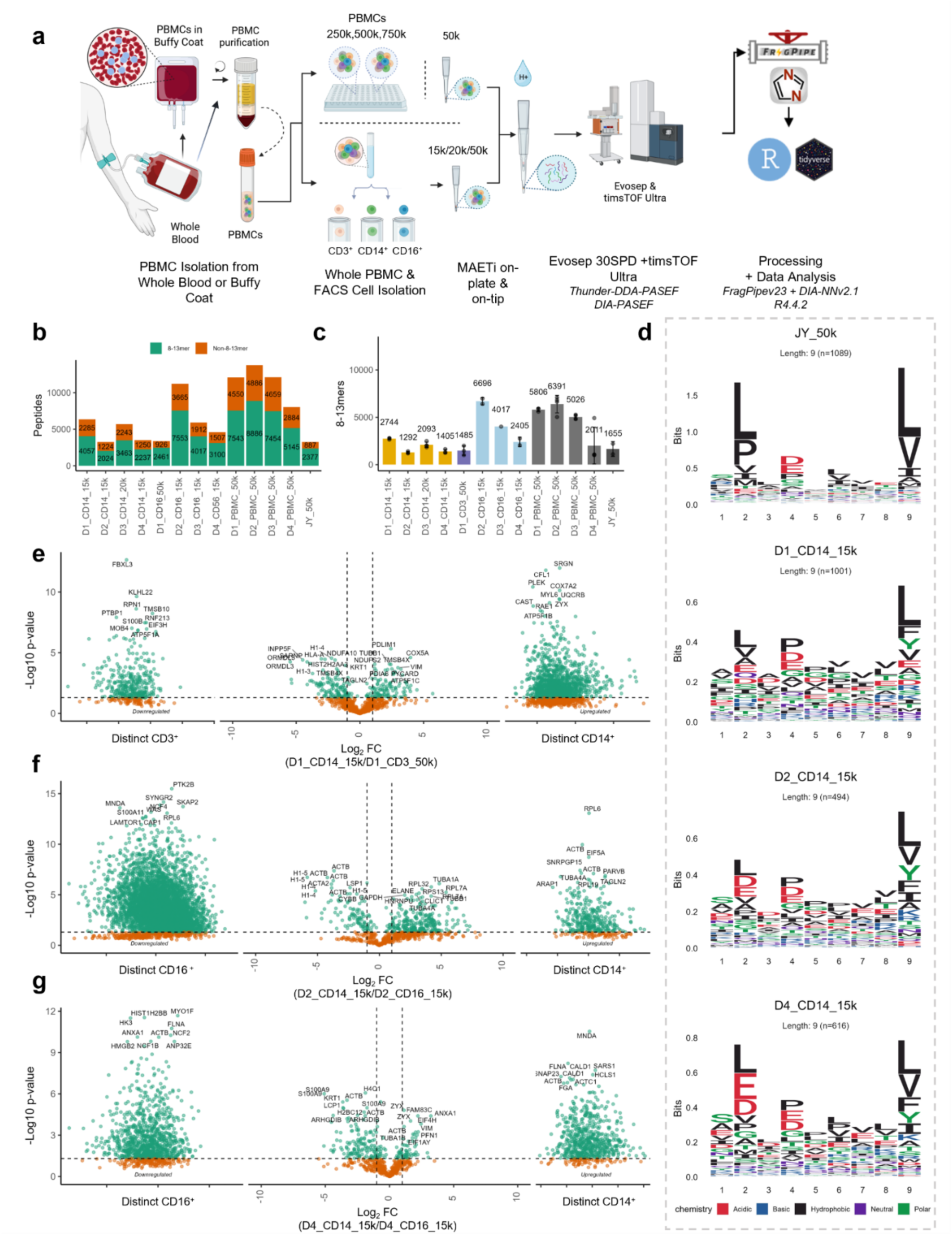
MAETi enables immunopeptidome profiling of fluorescence-activated sorted immune cells from human PBMCs. **a** Human peripheral mononuclear cells (PBMCs) of four healthy donors (D1-4) were purified and CD3^+^, CD14^+^ and CD16^+^CD56^+^ cells were isolated using fluorescence-activated cell sorting. Sorted cells were subjected to the MAETi on-tip format at 1.5E4 (15k) or 2E4 (20k) CD16^+^CD56^+^ or CD14^+^ cells (**bottom**) or 5E4 (50k) cells in case of CD3^+^ and the whole, unsorted PBMCs (**top**). Donor-specific spectral libraries were generated subjecting whole PBMCs at 25E4 (2.5E5; D1-4), 5E5 (5E5; D2-4)) and 75E4 (7.5E5, D1) cells to the MAETi on-plate format (top) using Thunder-ddaPASEF (n=3-4). All samples were pre-separated and measured on an Evosep (30SPD) coupled to a timsTOF Ultra. All samples of sorted cells were acquired in diaPASEF (n=1-4^#^ per cell type and donor). Raw files were processed in FragPipev23 with integrated DIA-NNv2.1 creating individual jobs per donor and generating spectral libraries on-the-fly for DIA-Quant analysis. Donor 1 (D1) was HLA-typed. *Created in BioRender. Tenzer, S. (2026)* https://BioRender.com/3ms0n1d. **b** Total number of peptides identified in diaPASEF (n=1-4) for 1.5E4 or 2E4 CD14^+^, 1.5E4 CD16^+^CD56^+^ or 5E4 CD3^+^ cells and 5E4 whole PBMCs, divided in 8-13mers and other peptide lengths. **c** Peptides with the expected size (8-13mers) identified on average per sample **d** Peptide sequence logos of 9mers identified for JY (control) and isolated cell types **e-g** Volcano plots with differentially (Log_2_FC > -10; Log_2_FC < 10) and distinct (Log_2_FC < -10; Log_2_FC > 10) 8-13mers comparing isolated cell types within donors; sub-panels at the sides represent exclusive identifications for each cell type. ^#^For label clarity thousands of cells (cells*1E3) were abbreviated with “k” and CD16^+^CD56^+^ cells were referred to as CD16^+^ cells. Technical replicates were as follows: 1.5E4 CD14^+^ cells n_[D1,D2,D4]_ = 3; 2E4 CD14^+^ cells n_[D3]_ = 4; 1.5E4 CD16^+^CD56^+^ cells n_[D2,D4]_ = 2 and n_[D3]_ = 1; 5E4 CD3^+^ cells n_[D1]_ = 3; 5E4 whole PBMCs n_[D1-D4]_ = 4; 5E4 JY n_JY_ = 2. Bars with error bars correspond to *average ± sd* with points indicating the individual values.

The DDA analysis of 2.5E5 to 7.5E5 PBMCs resulted in the identification of up to 12,596 unique 8-13mers. In the 3 to 4 replicates of 5E4 PBMCs acquired in DIA, 5,145 (D4, n=4) to 8,886 (D2, n=4) 8-13mers were identified. In the sorted cells, the number of 8-13mers detected varied across individuals but also cell types from the same donor (Fig. 6b, c). For instance, in D2, the sorted monocytes (CD14_15k, n=3) yielded 2,024 8-13mers in total and 1,292 ± 111 on average (± sd), while the NK-cells (CD56_15k, n=2) resulted in 7,553 8-13mers in total and 6,697 ± 385 on average (± sd). This ∼3.8- to 5-fold difference in the immunopeptidome coverage from the same donor and similar cell inputs already highlights the large differences in the MHC1 ligand repertoire across cell types. Nevertheless, the low standard deviations show the robustness of the workflow. Indicating potential MHC1ps, the proportion of 8-13mers detected in DIA for both PBMCs and sorted cells ranged from 60% to 74% (Supplementary Fig. 6b).

To verify the enrichment of potential MHC1ps via MAETi in healthy donors, we next evaluated sequence motifs obtained from the eluted ligands. Cells from D1 were HLA-typed, but due to technical reasons this was not possible for D2-4. For D1 (HLA-A*02, HLA-B*39, HLA-B*62/HLA-B*15), the enriched 8-13mers followed characteristic motifs exhibiting hydrophobic anchors at amino acid position 2 (P2) and position 9 (P9) for HLA*02, basic residues (H/R) at P2 for HLA-B*39 as well as polar tyrosine (Y) at P9 for HLA-B*62/HLA-B*15. Similarly, we evaluated the sequence motifs obtained for non-HLA-typed D2-4. Across both PBMCs and sorted cells, the enriched 8-13mers of D2 to D4 exhibited defined 9mer sequence motifs with anchor amino acids P2 or P9 as expected for MHC1ps (Fig. 6d; Supplementary FigS6d). Here, particularly the C-terminal anchor at P9 is dominated by hydrophobic amino acids (L, F, V, I), occasionally less dominant polar Y or basic K. At P2, the hydrophobic amino acid L or acidic amino acids D or E dominate the motifs. Moreover, supervised motif deconvolution via MHCMotifDecon (v1.0)^41^ indicates good correspondence with several HLA-supertypes (Supplementary Fig. S6d). For instance, HLA-A*02:01 association was found in all donors, which is not surprising given that it is one of the most common alleles in populations of European descent. Altogether, this shows that MAETi enables the identification of potential MHC1 ligands of unsorted and FACS-sorted cell populations even from primary human healthy donor samples.

To evaluate qualitative differences among sorted cell types within donors, we next investigated the overlap and uniqueness of 8-13mers detected in at least 60% of the replicates of each condition (Supplementary Fig. S6c). Qualitatively, all cell types showed some level of uniqueness and, while for D1 it was highest for CD14^+^ cells, for D2-4 it was always highest for CD16^+^CD56^+^ 8-13mers. In all cases, the overlap between the two cell types of interest ranged between 12.5% (D2) to 28.5% (D1). This indicates shared or unique potential immunopeptides present in the individual cell types.

To further evaluate the differential signatures between CD14^+^ and CD16^+^CD56^+^ cells or CD3^+^ cells within each donor, we integrated both qualitative and quantitative 8-13mer information obtained by the MAETi DIA (shown for D1, D2 and D4; Volcano Plots Fig. 6e-g). Irrespective of the cell types within donors, MAETi enabled the assessment of both differentially regulated and cell-type-specific 8-13mers. Although most 8-13mers stemmed from housekeeping proteins, others are reported to be involved or upregulated in immune system processes or describe activities of the particular target cell type (STRING^42^ Top20 Gene Ontology (GO) Terms Supplementary Fig. S7a- Fig. S7c). In fact, in some donors these immune signatures were already evident from the shared but differentially presented 8-13mers. For example, for the CD16^+^CD56^+^ cells of D4, the top 20 GO terms include “Cell killing”, “Leukocyte mediated cytotoxicity”, or “Regulation of inflammatory response”. Interestingly, for the CD3^+^ cells of D1 top20 GO terms of shared but differential 8-13mer-derived proteins include “Negative regulation cell killing” or either “Negative regulation natural killer cell mediated cytotoxicity” but at the same time “Positive regulation T cell mediated immunity” or “Positive regulation of T-cell mediated cytotoxicity” and “antigen processing and presentation” in the class I pathway.

These results demonstrate that MAETi enables immunopeptidome profiling of FACS-sorted cell populations from healthy human donors, providing both qualitative and quantitative insights into presented immunopeptidomes of sorted cell types with a minimal input of 1.5E4-5E4 cells.

## 3. Discussion

In the past years, the landscape of immunopeptidomics enrichment techniques was dominated by IP^4,16,43^. This is partly due to reports about yield and unspecific MHC1p elution using MAE^16,43^. However, more recently, Sturm *et al.* proved that an optimized MAE protocol can yield comparable and complementary results to bulk IP, with similar MHC1p enrichment specificity.^16^ Thus, we based our initial MAETi buffer on the protocol proposed by Sturm *et al.*^16^ and further improved it by replacing citric acid with β-alanine which resulted in lower background ions and interferences. This optimization improved both MHC1p coverage and reproducibility of our workflow. When scaled up to bulk sample preparation, the β-alanine-based MAE provided identification characteristics comparable to those of conventional IP.

Using two protocol layouts (on-tip or on-plate, Fig. 5a), MAETi is scalable for different input amounts and types of applications and can be applied from as low as 1E4 JY cells to up to 1E6 JY cells, enriching MHC1ps with high specificity. MAETi-prepared samples can readily be analyzed with both DDA and DIA approaches, enhancing its versatility and analytical depth. Spectral library-based diaPASEF improved coverage, dynamic range, and data completeness for MHC1p identifications and yielded on average 3,454 predicted binders per 1E4 JY cells and 6,296 predicted binders per 1E6 JY cells. Additionally, MAETi enabled robust detection of SAVs, establishing its rationale for potential neoantigen discovery for low-input immunopeptidomics.

One strength of MAETi is its minimal sample handling. Peptide adsorption on plastic surfaces during proteomics or peptidomics sample handling is known to substantially affect sample yield.^44–47^ In fact, several studies have reported substantial peptide losses during IP-based enrichments, which typically involve multiple sample-handling steps, including repeated washing steps after MHC-peptide complex capture^17–21^. While others have also adopted MAE strategies for low-input immunopeptidomics, their approach still requires ultrafiltration and off-line desalting, which can result in sample loses. For instance, after our first report of MAETi^48^, Bathini *et al*. ^49^ reported a method yielding 500 antigens from 100,000 cells. In comparison, due to minimal sample handling and therefore lower peptide losses, MAETi is not only compatible with lower sample input material but also substantially quicker, providing ready-to-analyze samples in less than 3 hours starting from washed cells, and higher coverage. In addition, acquired MAETi data from cryopreserved cells (see “MAETi enables MHC1 ligandomics of FACS-sorted immune cells”) demonstrate the utility and flexibility beyond in-culture applications.

In line with more efficient sample handling and minimized losses, in the past 4 years, IP-based microfluidic approaches have also been explored for low-input immunopeptidomics.^17,22,23^ In comparison to a microfluidic IP layout (“peptiCHIP”) by Feola *et al.* using JY cells^22^, MAETi provides enhanced MHC1p length profiles and higher 8-13mer proportions at equal (1E6) or lower input cell numbers. However, in contrast to microfluidics, MAETi does not require specialized equipment or expertise^17^, enabling higher throughput, and easy and quick implementation by other labs. This might be one reason, why other labs investigated alternative protocols based on MAE too.

One limitation of our study is the use of self-fabricated GF-Filter tips. Although the procedure of fabricating filter-tapered pipette tips has been described before by Rappsilber et.al.^50^, throughput and technical reproducibility of MAETi may be increased by replacing self-fabricated GF-Filter tips with commercially manufactured versions. Exploring automated workflows or integration with high-throughput systems could further expand the utility of MAETi, making it even more accessible for large-scale immunopeptidome studies. In addition, novel diaPASEF methods such as slicePASEF^51^ and midiaPASEF^52^ continue to push sensitivity and analytical depth. Coupled with novel instrument platforms and advanced processing solutions enabling directDIA searches (e.g., FragPipe v22.0^53^ or Spectronaut^54^) or DIA analysis using de novo predicted immunopeptide spectral libraries (e.g., PeptDeep/PeptDeep-HLA^55^), these technologies can further enhance the sensitivity of MAETi.

In the present study using MAETi, we profiled immunopeptidomes of 1.5E4 CD16^+^CD56^+^ and 2E4 CD14^+^ cells isolated from three different PBMC donors with unknown HLA-typing. Additionally, we included MAETi data obtained from 1.5E4 CD14^+^ and 5E4 CD3^+^ cells of a 2-digit HLA-typed donor. In all cases, isolated 8-13mers exhibited strong peptide sequence motifs with anchor amino acid residues mostly at P2 and P9 characteristic for MHC1ps. MAETi further enabled us to assess distinct and differential 8-13mer signatures for isolated cell types. This is particularly valuable since, even within the same cell type and HLA background, multiple factors shape immunopeptidomes both qualitatively (e.g., induction of the immunoproteasome) and quantitatively (e.g., HLA expression, PTMs, and protein turnover).

Although the acquisition of immunopeptidomes from magnetic-sorted cell populations has been shown before*^7^*, to our knowledge, immunopeptidomics of flow cytometry-sorted cells based on multiple lineage markers (HLA-A, -B, -C^+^ and CD3^+^, CD14^+^ as well as CD16^+^CD56^+^) in the range of 1.5E4-5E4 cells has not been reported yet. Marino *et al.^7^*analyzed immunopeptidomes of magnetically sorted CD14^+^, CD4^+^, CD8^+^, and CD19^+^ cells obtained by immunoprecipitation of 5E6 to 17E6 cells undergoing activation. In contrast, MAETi enabled scale-down of required input amounts 100-fold for CD3^+^ and 300-fold for CD14^+^ cells, facilitating targeting of other rare cell subsets of interest.

The optimized MAETi protocol introduced in this study offers a cost-effective and scalable approach for MHC-I peptide enrichment, achieving results comparable to IP. At the same time, low input requirements and efficient sample handling enable immunopeptidome analysis from as few as 1E4 JY cells or sorted immune-cell populations. This could be particularly valuable for investigating heterogeneity and dynamics in antigen presentation, both qualitatively and quantitatively. Moreover, we anticipate that MAETi will enable immunopeptidomics experiments requiring high sensitivity and throughput across various other biological applications, such as drug screenings and kinetics. Thus, MAETi holds significant potential for advancing tumor immunology, and informing personalized immunotherapy and drug discovery.

## 4. Materials and Methods

### 4.1. Patient material

Fully anonymized healthy patient whole blood and buffy coats were obtained via the Transfusion Medicine Department of the University Medical Center Mainz in accordance with the Declaration of Helsinki and institutional guidelines and ethical regulations for research use. All biospecimens including whole blood, buffy coats and isolated PBMCs were kept at room temperature (RT).

### 4.2. Materials and substances

Analytical grade substances and reagents were used for sample preparation, and were acquired from Sigma Aldrich (Merck), unless otherwise stated. Water, acetonitrile, and formic acid were LC-MS grade products acquired from Carl Roth. LoBind tubes (Eppendorf) were employed to minimize sample loss.

### 4.3. Cell culture

The human B lymphoblastoid cell line JY (CVCL_0108) was purchased from ATCC and the human Burkitt lymphoma cell line Raji (CVCL_0511) was obtained from the DSMZ-German Collection of Microorganisms and Cell Cultures. Both cell lines were maintained in RPMI1640 medium supplemented with 10 % FCS (Gibco (v/v)), 2 mM glutamine, 1 mM sodium pyruvate, 100 units/ml penicillin, and 100 μg/ml streptomycin. Suspension cell cultures were harvested at 220 x g for 10min and washed three times with 1x PBS at 4°C prior counting and either submitting to MAE as whole cells or freezing at −80 °C until further use for IP.

### 4.4. Isolation and Purification of PBMCs

For each donor, buffy coat was diluted 1:2 with PBS (RT) before carefully layering 30 mL diluted buffy coat on top of 20 mL Ficoll Paque medium (Cytiva™ Ficoll Paque PREMIUM, 17544202) at RT. Centrifugation at 400 x g without brake for 30 min at RT resulted in a layer of PBMCs which were collected using a pipette and added to conical tube pre-filled with 3 times volume of PBS + 0.2%BSA. Supernatant was removed after centrifugation at 500 x g for 15 min at RT, and washing steps were repeated twice. In the last washing step, cells were resuspended in PBS + 0.2%BSA and counted with a Neubauer counting chamber using trypan blue after straining through a 40 µm cell strainer (Greiner Bio-One, Cell strainer EASYstrainer™, Art. Nb.: 542040) and centrifuged at 300 x g for 5 min at RT. After careful removal of supernatant, cell pellet was resuspended in cryomedium (10% DMSO in FCS) at a concentration of 4×10^6^ cells/mL. Cells were frozen at a controlled rate of -1°C per minute using a freezing container placed in a -80°C freezer. For long-term preservation vials should be transferred to liquid nitrogen storage.

### 4.5. Fluorescence-activated cell sorting (FACS) of PBMCs

Cryo-preserved cells were quickly thawed in a water bath at 37°C. Cells were immediately washed twice with prewarmed PBS + 0.2%BSA and supernatant was removed by centrifugation at 300 x g for 5 min at RT. For cell staining, cells were incubated in PBS/0.2%BSA supplemented with anti-Fc Receptors (Human TruStain FcX™; BioLegend®, Cat. Nb.: 422301) and Monocyte blocker (True-Stain Monocyte Blocker™; BioLegend®, Cat. Nb.: 426103) for 20 min at RT to prevent non-specific binding of staining antibody. Then cells were stained for 20 min at RT in the dark with anti-CD16 RB744 (clone 3G8; BD Biosciences), anti-CD14 APC H7 (clone SMФP9; BD Biosciences), anti-CD3 BV480 (clone UCHT1; BD Biosciences), HLA-ABC BV788 (clone G46-2.6; BD Biosciences) and CD56 BUV737 (clone NCAM16.2; BD Biosciences). To exclude dead cells, 0.1 µg/ ml 4′,6-diamino-2-phenylindole (DAPI, Roche) was utilized.

Sorting was performed on a BD FACSAria™ II (BD Biosciences) using a 70 µm nozzle. Laser powers and detection filters were configured according to the manufacturer’s specifications. For the lymphocyte gating, forward and side scatter area (FSC-A vs. SSC-A) was used, and single cells were defined by FSC-H vs. FSC-A. HLA-ABC^+^ cells were gated prior to sorting of CD3^+^, CD14^+^ and CD16^+^CD56^+^ cells. Sorted cells were collected into Protein LoBind tubes and the stream was aligned slightly off-center to reduce shear stress. Cells were stored on ice before the subsequent MAETi procedure. Exemplarily, the flow cytometric identification of HLA-ABC^+^ & CD14^+^, CD16^+^CD56^+^, CD3^+^ cells is shown in Supplementary Fig. S5.

### 4.6. MAE Filter Tip (MAETi) Preparation

Glass fiber filter tips were prepared as described by Rappsilber *et al.*^50^ using a blunt-cut injection needle (BD microlance 3, 1.2 x 40 mm, 18G) and a 1.7 µm glass fiber (GF) membrane (GE Healthcare, Whatman^TM^ CAT No. 1820-030) tapered into a Protein LoBind tip (Greiner BioOne Sapphire Tips). Prior utilization in the MAETi procedure (Section 4.7), fabricated tips were washed twice with PBS and once with MAE buffer at 220 x g, for 1 min.

Evotips were placed in isopropanol and washed twice with 50 µL ACN + 0.1% FA and once with 50 µL H_2_O + 0.1% FA before placing the Evotips in 0.1% FA (v/v) in Water, following one more washing and equilibration step with 50 µL 0.1% FA (v/v) in Water. After washing, conditioning and equilibration, Evotips were topped-up with 70 µL 0.1% FA (v/v) in Water and stored at 4°C prior sample loading (Section 4.7).

### 4.7. Immunopeptide enrichment by mild acid elution in a tip (MAETi)

All next steps were either performed at 4°C or on ice to ensure sample integrity. Depending on the input cell number, we used two protocol layouts (also see Fig. 5a) to profile input amounts across a range from 10,000 (1E4) to 1,000,000 (1E6) cells.

In brief, cells were washed and prepared according to their input number, transferred to the respective protocol layout (on-Tip or on-Plate, Fig. 5a), followed by MAE buffer treatment and peptide capture on the Evotip. For both recommended protocol versions, eluted peptides captured by the Evotip were subsequently washed once with 50 µL 0.1% FA (v/v) in Water and finally topped up with 200 µL 0.1% FA (v/v) in Water.

The original citric acid-based MAE buffer formulation was adapted from Sturm *et al.*^16^ adjusted to pH 3.3. Note, that for Section 2.2 the citric acid- and a β-alanine-based MAE buffer were compared but used at equimolar concentrations. Pure substances were compared using equimolar amounts of pure buffer substances prepared in glass bottles, directly transferred to the Evotip.

### 4.8. Immunopeptide enrichment by bulk mild acid elution (MAE)

The initial mild acid elution (MAE) protocol and buffer recipe was based on a protocol published by Sturm *et al.*^16^, with slight adaptations. Throughout the MAE procedure, all steps were performed at 4°C or on ice. Briefly, pre-washed and counted whole cells (see Section 2.3) were pelleted at 220 x g, 10 min and 40E6 cells per replicate (n=3) were resuspended in ice-cold MAE buffer (pH 3.3) adapted from Sturm *et al.*^16^ by pipetting up and down three times and gently tapping the tubes during a 1 min incubation on ice. After incubation, the MAE supernatant was immediately removed by centrifugation at 300 x g for 5 min and further cleared by centrifugation at 350 x g for 10 min and 3,500 x g for 15 min. To get rid of potential cellular debris, retrieved MAE supernatant was further cleared twice at 16,000 x g for 20 min, transferring the supernatant to a new tube in between.

### 4.9. Immunopeptide enrichment by bulk immunoprecipitation (IP)

Class I MHC immunoprecipitation was performed as previously described by Gomez-Zepeda *et al*.^27^ (detailed protocol in Gomez-Zepeda *et al.* 2024^56^). Briefly, frozen aliquoted and PBS-washed cell pellets at 40E6 cells per replicate (n=3) were thawed in lysis buffer (1% CHAPS (v/v) in PBS), subjected to sonication (Diagenode™ Bioruptor Plus: 10 cycles, 30s on/off for a total duration of 10 min) and incubated overnight at 4°C with W6/32 antibody (Leinco™ Anti-Human HLA-A, B, C-purified in vivo GOLD functional grade, P/N H263) coupled to CNBr-activated Sepharose (Cytiva™; P/N 17098101). Captured MHC class I peptide complexes were washed with PBS, MS-grade H_2_O and eluted with 0.2% trifluoroacetic acid (TFA, (v/v)).

### 4.10. Ultrafiltration and peptide desalting of bulk IP and MAE samples

Samples prepared by bulk IP and bulk MAE were ultrafiltered (Sartorius® Vivacon 500, 10.000 MWCO Hydrosart) prior desalting by SPE on a Hydrophilic-Lipophilic-Balanced (HLB) sorbent (Oasis HLB 96-well μElution Plate, 2 mg Sorbent per Well, 30 μm, Waters Corp.). Peptides were eluted from the HLB resin with 35% ACN (v/v) in MS-grade H_2_O + 0.1% TFA (v/v), lyophilized and reconstituted in 15 µL 0.1% FA (v/v) in MS-grade H_2_O prior LC-MS analysis (Section 4.11). Samples were injected utilizing approximately 1 µL (∼ 50 ng; ∼ 2 million cell equivalents (MCEs)) of each replicate in technical triplicates.

### 4.11. Bulk IP and MAE LC-MS acquisitions on timsTOF Pro 2

Samples for the bulk comparison (Section 2.3; Fig. 4) were separated using a NanoElute 2 LC system coupled to a timsTOF Pro 2 mass spectrometer. Desalted peptides obtained from bulk-MAE and bulk IP preparations (Section 4.8 to 4.10) were directly injected onto a C18 reversed-phase (RP) Aurora 25 cm analytical column (25 cm x 75 µm ID, 120 Å pore size, 1.7 µm particle size, IonOpticks, Australia) and separated using gradients increasing the proportion of phase B (ACN + 0.1% FA (v/v)) to phase A (water + 0.1% FA (v/v)) over 47 min. A Captive Spray source was used for ionization, with a capillary voltage of 1600 V, dry gas at 3.0 L/min, dry temperature at 180 °C, and TIMS-in pressure of 2.7 mBar. Bulk IP and MAE samples (as obtained under Section 4.8 to 4.10) were analysed in three injection replicates for each biological/preparation replicate, using a target of 50 ng (∼2 MCEs) per injection (detailed outline in Supplemental) acquiring data utilizing a non-MHC-tailored “stepped” ddaPASEF precursor selection polygon including singly-charged ions for unbiased comparison.

### 4.12. MAETi LC-MS acquisition on timsTOF SCP

MAETi-eluted peptides captured on Evotips in Section 4.7 were separated at 50 °C on a C18 RP Aurora Elite 25 cm analytical column (15 cm x 75 µm ID, 120 Å pore size, 1.7 µm particle size, IonOpticks, Australia) using the 40SPD whisper method on the Evosep One nanoLC system coupled to a timsTOF SCP. A Captive Spray source was used for ionization, with a capillary voltage of 1400 V, dry gas at 3.0 L/min, dry temperature at 200 °C, and TIMS-in pressure of 2.7 mBar. For proof of principle experiments (Section 2.1) 5E4 cell samples were injected utilizing a non-MHC-tailored “stepped” polygon and the MHC-tailored “thunder” polygon^27^ (Fig. 3 2a, b). Buffer comparisons (Section 2.2) were acquired applying the same settings using the “stepped” polygon for unbiased comparison. All acquisitions on the timsTOF SCP utilized a 100 ms TIMS ramp, a scan range of from *m/z* 100-1700, 10 PASEF ramps and a cycle time of 1.17 sec.

### 4.13. MAETi LC-MS acquisition on timsTOF Ultra

MAETi-eluted peptides captured on Evotips in Section 4.7 were separated at 40 °C on a C18 Reversed-phase (RP) analytical EV1137 Performance column (ReproSil-Pur C18, 1.5 µm beads by Dr Maisch. 15 cm x 150 µm) using the 30SPD method on the Evosep nanoLC system and 20 µm zero-dead volume CaptiveSpray 2 Emitters (Bruker, Part No:1811112). An Ultra source was used for ionization, with a capillary voltage of 1500 V, dry gas at 3.0 L/min, dry temperature at 200 °C at 1 mBar Funnel and 2.38 mBar TIMS-in pressure. Ions were accumulated and separated utilizing a 60 ms TIMS ramp. Acquisitions in Thunder-ddaPASEF utilized the “thunder” MHC-tailored precursor isolation polygon^27^ selecting precursors (charge +1 to +3) in the range of *m/z* 100-1700 and 0.65 to 1.76 V*s/cm^2^ for fragmentation utilizing 16 PASEF ramps per cycle, resulting in total cycle time of 1.12 sec. Precursors were isolated with 2 Th windows below *m/z* 700 and 3 Th above and, when reaching a target intensity of 2E4 and an intensity threshold of 500 arbitrary units, actively excluded for 0.4 min. High-sensitivity mode was activated and denoising in the data processing step was switched off.

### 4.14. JY whole exome sequencing

Database search was conducted on a customized database (fasta file) obtained from whole exome sequencing (WES) of JY.

In brief, genomic DNA was extracted from 6,000,000 JY cells using the cell-lysate homogenizer QIAshredder (QIAGEN, Cat# 79656) and AllPrep DNA/RNA Mini kit (QIAGEN, Cat# 80204). Whole-exome sequencing libraries were prepared using the SureSelect XT HS V2 DNA system (Agilent Technologies) in combination with the SureSelect Human All Exon V7 capture kit according to the manufacturer’s protocol. Libraries were sequenced with the Illumina NovaSeq 6K PE 100 S1 technology.

### 4.15. Construction of the personalized database from WES data

The JY custom database containing its amino acid deviation from the reference proteome database was generated using the whole exome sequencing data. The fastq data was quality controlled using fastqc (v0.12.0)^57^ and pairwise aligned on the human reference genome GRCh38 using Bowtie2 (v2.5.2)^58^ with the “fast-local” preset. Alignment output is converted to bam file via samtools (v1.18)^59^ and subsequently prepared for variant calling using “fixmate”, “sort”, “index”, “markdup”. Variant calling was done using bcftools (v1.18) “mpileup” command. Position of variants from the resulting VCF file are mapped to transcripts using GENCODE^60^ hg38.knownGene.gtf as reference. The absolute genomic positions are converted to transcript relative positions using AnnotationHub (v3.9.0)^61^ library. Transcripts are “mutated” accordingly while keeping multiple variations of the same transcript in case of heterozygosity. Transcripts are then translated and reference proteins are replaced by its related mutated translated transcripts.

### 4.16. Peptidomics database search

For method optimization, data analysis was performed in PEAKS XPro (v10.6, build 20201221). The protein database was composed of the UniProtKB (Swiss-Prot) reference proteomes of Homo sapiens (Taxon ID 9606, downloaded 02/Feb/2020), Epstein-Barr virus (strain GD1, Taxon ID 10376, downloaded 06/Feb./2022), supplemented with a list of 172 possible contaminants. For database searches, protein in silico digestion was configured to unspecific cleavage and no enzyme. Up to two variable modifications per peptide were allowed, including Methionine oxidation, cysteine cysteinylation, and Protein N-terminal acetylation. The option timstof_feature_min_charge (in file PEAKSStudioXpro\algorithmpara\feature_detection_para.properties) was set to 1 to allow the identification of singly-charged features. Raw LC-MS files were loaded with the configuration for timsTOF ddaPASEF data with CID fragmentation. Mass tolerance was set to 15 ppm for MS1 and 0.03 Da for MS2.

For the titration experiment with JY cells (Section 2.4) at different cell inputs data were analysed using Fragpipe (FragPipe v22.0 + integrated DIA-NN v1.8.2 beta 8; MSFragger version 4.1, IonQuant version 1.10.27, Percolator version 3.6.5, Philosopher version 5.1.1) applying the default “Nonspecific-HLA-DIA" workflow. The WES-based personalized database as generated under section 4.14 was complemented with 172 common LC-MS contaminants and the genome of EBV-strain B95-8 (2024-12-18-decoys-JY1_WES_24407_20586_conts_2023-6-29_EBV_B95-8.fasta).

For MAETi of FACS-sorted cells (Section 2.5), data were analysed using Fragpipe (FragPipe v23.0 + integrated DIA-NN version 2.1.0 Academia; DIA-Umpire version 2.3.2, diaTracer version 1.2.5, MSFragger version 4.2, MSBooster version 1.3.9, IonQuant version 1.11.9, Philosopher version 5.1.1) applying the default “Nonspecific-HLA-diaPASEF" workflow, creating donor-specific analyses via batch processing. Fasta database was based on UniProt release 2020-3-2 complemented with 172 common LC-MS contaminants the genome of EBV-strain B95-8 as well as contaminant murine Ig gamma-1 chain “mIGHG1” (2025-05-14-decoys-human_20365_conts_172_2020-3-2_2-0-11_mIGHG1_EBVB95.fasta).

In both cases, DDA files were loaded along the DIA files for on-the-fly spectral library generation, selecting “DIA-Quant” for the DIA files. Applied FragPipe versions are noted within the sections. For both FragPipe processing runs, fasta and log files are supplied along the JPost submission^62–64^.

### 4.17. Experiment design

Experimental descriptions including number of replicates per conditions are detailed in the respective result sections.

### 4.18. Data analysis and statistics

Data analysis, statistical analysis and visualization were done mostly using in house R scripts^65^. The main R packages used were as follows; the statistical difference was assessed by two-sided t-test using ggpubr (v. 0.4.0)^66^; plots were generated using ggplot2 (v. 3.4.0)^66^; Venn plots with ggvenn (v. 0.1.9)^67^; upset plots with ggupset (v. 0.3.0)^68^; and sequence logos were generated using ggseqlogo^69^. OpenAI’s ChatGPT^70^ (GPT-4 & GPT-5) was employed strictly as a software tool to generate and troubleshoot code snippets. Its use was limited to generic coding tasks, with no input of study data or involvement in the scientific reasoning of this work.

Note that throughout the manuscript “Peptides” refers to peptides including their modifications, “Sequences” refers to the “stripped peptide” (no modifications) and binders refers to “Sequences” predicted to be weak or strong MHC binders for the sample-specific HLA alleles. Binding prediction was performed using NetMHCpan-4.130 and sequence clustering with GibbsCluster-2.064, both via MhcVizPipe (v0.7.9)^28^. The MHC selected allotypes used for prediction were as follows: JY (HLA-A*0201, -B*0702, -C*0702), Raji (HLA-A*03:01, -B*15:11, -C*03:04, -C*04:01).

For differential analyses, data were filtered by length (8-13mers) and data completeness (IDRate) i.e., how often a peptide was detected across multiple replicates. To assess differential abundance between conditions A and B, log₂ fold changes (FCs) were calculated as:

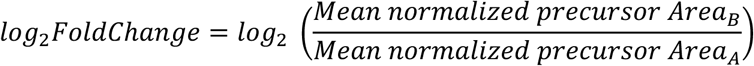

To evaluate statistical significance, two-sample t-tests were performed assuming equal variances and independent samples. The t-statistic was computed as:

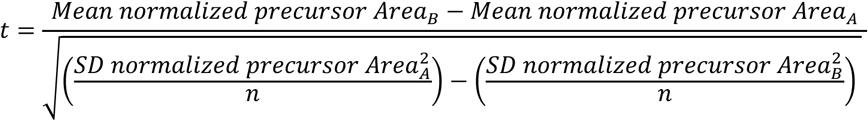

where n is the number of biological replicates per condition (e.g., n=4). Degrees of freedom were set to 2(n − 1) = 6. Two-tailed p-values were obtained from the corresponding t-distribution. Significance threshold for up- or downregulated peptides were log2FC ± 1 and a p-value threshold of 0.05. To plot detections exclusive (“distinct”) to either condition A or B, peptides not found in any of the replicates of the other condition were assigned a very small value (1e-10) resulting in extreme log2 FC values separating unique from differential identifications in the volcano plots.

### 4.19. Data availability

The mass spectrometry immunopeptidomics will be deposited to the ProteomeXchange Consortium via jPOST^62–64,71^ upon publication.

### 4.20. Code availability

All evaluations were performed with in-house developed code.

## 6. Acknowledgements

We acknowledge the support of the Immunopeptidomics Platform of HI-TRON Mainz. We would like to acknowledge Lucas Kleinort (HI-TRON, Mainz, Germany) for his technical assistance. We would like to thank our colleagues, Dr. Malte Sielaff and Marian Scherer for their invaluable instrument support throughout this study. Figures 1, 2a, 3a, 4a, 5a, and 6a were created using BioRender in accordance with their licensing agreement.

## 7. Author contributions

Conceptualization: DGZ, ST. Methodology: JB, DGZ, YY, ST. Software: JB, YC. Validation: JB. Formal analysis: JB. Investigation: JB, YY, MF, AA, RP. Resources: DGZ, CC, UD, ST, RP. Data Curation: JB, DGZ. Writing - Original Draft: JB, DGZ. Writing - Review & Editing: all the co-authors. Visualization: JB, DGZ. Supervision: DGZ, ST. Project administration: DGZ, ST. Funding acquisition: ST

## References

1. Bassani-Sternberg, M. Mass Spectrometry Based Immunopeptidomics for the Discovery of Cancer Neoantigens. in 209–221 (2018). doi:10.1007/978-1-4939-7537-2_14.

2. Nelde, A. et al. Immunopeptidomics-Guided Warehouse Design for Peptide-Based Immunotherapy in Chronic Lymphocytic Leukemia. Front Immunol 12, (2021).

3. Fritsche, J. et al. Translating Immunopeptidomics to Immunotherapy-Decision-Making for Patient and Personalized Target Selection. Proteomics 18, (2018).

4. Purcell, A. W., Ramarathinam, S. H. & Ternette, N. Mass spectrometry–based identification of MHC-bound peptides for immunopeptidomics. Nat Protoc 14, 1687–1707 (2019).

5. Caron, E., Aebersold, R., Banaei-Esfahani, A., Chong, C. & Bassani-Sternberg, M. A Case for a Human Immuno-Peptidome Project Consortium. Immunity 47, 203–208 (2017).

6. Vizcaíno, J. A. et al. The Human Immunopeptidome Project: A Roadmap to Predict and Treat Immune Diseases. Molecular & Cellular Proteomics 19, 31–49 (2020).

7. Marino, F. et al. Biogenesis of HLA Ligand Presentation in Immune Cells Upon Activation Reveals Changes in Peptide Length Preference. Front Immunol 11, (2020).

8. Chong, C. et al. High-throughput and Sensitive Immunopeptidomics Platform Reveals Profound Interferonγ-Mediated Remodeling of the Human Leukocyte Antigen (HLA) Ligandome. Molecular & Cellular Proteomics 17, 533–548 (2018).

9. Bernhardt, M. et al. SILAC-based quantification reveals modulation of the immunopeptidome in BRAF and MEK inhibitor sensitive and resistant melanoma cells. Front Immunol 15, (2025).

10. Murphy, J. P. et al. Therapy-Induced MHC I Ligands Shape Neo-Antitumor CD8 T Cell Responses during Oncolytic Virus-Based Cancer Immunotherapy. J Proteome Res 18, 2666–2675 (2019).

11. Stopfer, L. E., Mesfin, J. M., Joughin, B. A., Lauffenburger, D. A. & White, F. M. Multiplexed relative and absolute quantitative immunopeptidomics reveals MHC I repertoire alterations induced by CDK4/6 inhibition. Nat Commun 11, 2760 (2020).

12. Murphy, J. P. et al. Multiplexed Relative Quantitation with Isobaric Tagging Mass Spectrometry Reveals Class I Major Histocompatibility Complex Ligand Dynamics in Response to Doxorubicin. Anal Chem 91, 5106–5115 (2019).

13. Nelde, A., Kowalewski, D. J. & Stevanović, S. Purification and Identification of Naturally Presented MHC Class I and II Ligands. in 123–136 (2019). doi:10.1007/978-1-4939-9450-2_10.

14. Wahle, M. et al. IMBAS-MS Discovers Organ-Specific HLA Peptide Patterns in Plasma. Molecular & Cellular Proteomics 23, 100689 (2024).

15. Shunji Sugawara, Toru Abo & Katsuo Kumagai. A simple method to eliminate the antigenicity of surface class I MHC molecules from the membrane of viable cells by acid treatment at pH 3. J Immunol Methods 100, 83–90 (1987).

16. Sturm, T. et al. Mild Acid Elution and MHC Immunoaffinity Chromatography Reveal Similar Albeit Not Identical Profiles of the HLA Class I Immunopeptidome. J Proteome Res 20, 289–304 (2021).

17. Stutzmann, C. et al. Unlocking the potential of microfluidics in mass spectrometry-based immunopeptidomics for tumor antigen discovery. Cell Reports Methods 3, 100511 (2023).

18. Stopfer, L. E. et al. Absolute quantification of tumor antigens using embedded MHC-I isotopologue calibrants. Proceedings of the National Academy of Sciences 118, (2021).

19. Stopfer, L. E., D’Souza, A. D. & White, F. M. 1,2,3, MHC: a review of mass-spectrometry-based immunopeptidomics methods for relative and absolute quantification of pMHCs. Immuno-Oncology and Technology 11, 100042 (2021).

20. Hassan, C. et al. Accurate quantitation of MHC-bound peptides by application of isotopically labeled peptide MHC complexes. J Proteomics 109, 240–244 (2014).

21. Bijen, H. M. et al. Specific T Cell Responses against Minor Histocompatibility Antigens Cannot Generally Be Explained by Absence of Their Allelic Counterparts on the Cell Surface. Proteomics 18, (2018).

22. Feola, S. et al. PeptiCHIP: A Microfluidic Platform for Tumor Antigen Landscape Identification. ACS Nano 15, 15992–16010 (2021).

23. Li, X. et al. A microfluidics-enabled automated workflow of sample preparation for MS-based immunopeptidomics. Cell Reports Methods 3, 100479 (2023).

24. Purcell, A. W., Ramarathinam, S. H. & Ternette, N. Mass spectrometry–based identification of MHC-bound peptides for immunopeptidomics. Nat Protoc 14, 1687–1707 (2019).

25. Sturm, T. et al. Mild Acid Elution and MHC Immunoaffinity Chromatography Reveal Similar Albeit Not Identical Profiles of the HLA Class I Immunopeptidome. J Proteome Res 20, 289–304 (2021).

26. Nelde, A., Kowalewski, D. J. & Stevanović, S. Purification and Identification of Naturally Presented MHC Class I and II Ligands. in 123–136 (2019). doi:10.1007/978-1-4939-9450-2_10.

27. Gomez-Zepeda, D. et al. Thunder-DDA-PASEF enables high-coverage immunopeptidomics and is boosted by MS2Rescore with MS2PIP timsTOF fragmentation prediction model. Nat Commun 15, 2288 (2024).

28. Kovalchik, K. A. et al. MhcVizPipe: A Quality Control Software for Rapid Assessment of Small- to Large-Scale Immunopeptidome Datasets. Molecular & Cellular Proteomics 21, 100178 (2022).

29. Reynisson, B., Alvarez, B., Paul, S., Peters, B. & Nielsen, M. NetMHCpan-4.1 and NetMHCIIpan-4.0: improved predictions of MHC antigen presentation by concurrent motif deconvolution and integration of MS MHC eluted ligand data. Nucleic Acids Res 48, W449–W454 (2020).

30. Andreatta, M., Alvarez, B. & Nielsen, M. GibbsCluster: unsupervised clustering and alignment of peptide sequences. Nucleic Acids Res 45, W458–W463 (2017).

31. Skowronek, P., Voytik, J. & Willems, S. py_diAID. Preprint at https://github.com/MannLabs/pydiaid.

32. Skowronek, P. et al. Rapid and In-Depth Coverage of the (Phospho-)Proteome With Deep Libraries and Optimal Window Design for dia-PASEF. Mol Cell Proteomics 21, (2022).

33. Yu, F. et al. Analysis of DIA proteomics data using MSFragger-DIA and FragPipe computational platform. Nat Commun 14, 4154 (2023).

34. Tsou, C.-C. et al. DIA-Umpire: comprehensive computational framework for data-independent acquisition proteomics. Nat Methods 12, 258–264 (2015).

35. Kong, A. T., Leprevost, F. V, Avtonomov, D. M., Mellacheruvu, D. & Nesvizhskii, A. I. MSFragger: ultrafast and comprehensive peptide identification in mass spectrometry–based proteomics. Nat Methods 14, 513–520 (2017).

36. Yu, F. et al. Fast Quantitative Analysis of timsTOF PASEF Data with MSFragger and IonQuant. Molecular & Cellular Proteomics 19, 1575–1585 (2020).

37. Demichev, V., Messner, C. B., Vernardis, S. I., Lilley, K. S. & Ralser, M. DIA-NN: neural networks and interference correction enable deep proteome coverage in high throughput. Nat Methods 17, 41–44 (2020).

38. Demichev, V. et al. dia-PASEF data analysis using FragPipe and DIA-NN for deep proteomics of low sample amounts. Nat Commun 13, 3944 (2022).

39. Oliinyk, D. et al. diaPASEF Analysis for HLA-I Peptides Enables Quantification of Common Cancer Neoantigens. Molecular & Cellular Proteomics 24, 100938 (2025).

40. Li, K., Teo, G. C., Yang, K. L., Yu, F. & Nesvizhskii, A. I. diaTracer enables spectrum-centric analysis of diaPASEF proteomics data. Nat Commun 16, 95 (2025).

41. Kaabinejadian, S. et al. Accurate MHC Motif Deconvolution of Immunopeptidomics Data Reveals a Significant Contribution of DRB3, 4 and 5 to the Total DR Immunopeptidome. Front Immunol 13, (2022).

42. Szklarczyk, D. et al. STRING v11: protein-protein association networks with increased coverage, supporting functional discovery in genome-wide experimental datasets. Nucleic Acids Res 47, D607–D613 (2019).

43. Kuznetsov, A., Voronina, A., Govorun, V. & Arapidi, G. Critical Review of Existing MHC I Immunopeptidome Isolation Methods. Molecules 25, 5409 (2020).

44. Weikart, C. M., Breeland, A. P., Taha, A. H. & Maurer, B. R. Enhanced Recovery of Low Concentration Protein and Peptide Solutions On Ultra-Low Binding Microplates. Future Sci OA 5, (2019).

45. Rabe, M., Verdes, D. & Seeger, S. Understanding protein adsorption phenomena at solid surfaces. Adv Colloid Interface Sci 162, 87–106 (2011).

46. Kristensen, K., Henriksen, J. R. & Andresen, T. L. Adsorption of Cationic Peptides to Solid Surfaces of Glass and Plastic. PLoS One 10, e0122419 (2015).

47. Goebel-Stengel, M., Stengel, A., Taché, Y. & Reeve, J. R. The importance of using the optimal plasticware and glassware in studies involving peptides. Anal Biochem 414, 38–46 (2011).

48. Bathini, M. et al. MHC1-TIP enables single-tube multimodal immunopeptidome profiling and uncovers intratumoral heterogeneity in antigen presentation. Preprint at 10.1101/2025.07.17.664894 (2025).

49. Rappsilber, J., Mann, M. & Ishihama, Y. Protocol for micro-purification, enrichment, pre-fractionation and storage of peptides for proteomics using StageTips. Nat Protoc 2, 1896–1906 (2007).

50. Szyrwiel, L., Sinn, L., Ralser, M. & Demichev, V. Slice-PASEF: fragmenting all ions for maximum sensitivity in proteomics. bioRxiv 10.1101/2022.10.31.514544 (2022) doi:10.1101/2022.10.31.514544.

51. Distler, U. et al. midiaPASEF maximizes information content in data-independent acquisition proteomics. bioRxiv 10.1101/2023.01.30.526204 (2023) doi:10.1101/2023.01.30.526204.

52. Li, K., Teo, G. C., Yang, K. L., Yu, F. & Nesvizhskii, A. I. diaTracer enables spectrum-centric analysis of diaPASEF proteomics data. bioRxiv 10.1101/2024.05.25.595875 (2024) doi:10.1101/2024.05.25.595875.

53. Biognosys AG. Spectronaut 17. Preprint at (2022).

54. Zeng, W.-F. et al. AlphaPeptDeep: a modular deep learning framework to predict peptide properties for proteomics. Nat Commun 13, 7238 (2022).

55. Gomez-Zepeda, D., et al. High-coverage immunopeptidomics using timsTOF mass spectrometers with Thunder-DDA-PASEF boosted by MS2Rescore. Preprint at 10.21203/rs.3.rs-4849156/v1 (2024).

56. Andrews, S. FastQC A Quality Control tool for High Throughput Sequence Data.. Preprint at https://www.bioinformatics.babraham.ac.uk/projects/fastqc/ (2010).

57. Langmead, B. & Salzberg, S. L. Fast gapped-read alignment with Bowtie 2. Nat Methods 9, 357–359 (2012).

58. Danecek, P. et al. Twelve years of SAMtools and BCFtools. Gigascience 10, (2021).

59. Frankish, A. et al. GENCODE: reference annotation for the human and mouse genomes in 2023. Nucleic Acids Res 51, D942–D949 (2023).

60. Morgan, M. & Shepherd, L. AnnotationHub: Client to access AnnotationHub resources.. Preprint at doi:10.18129/B9.bioc.AnnotationHub (2025).

61. Vizcaíno, J. A. et al. ProteomeXchange provides globally coordinated proteomics data submission and dissemination. Nat Biotechnol 32, 223–226 (2014).

62. Okuda, S. et al. jPOSTrepo: an international standard data repository for proteomes. Nucleic Acids Res 45, D1107–D1111 (2017).

63. R Foundation for Statistical Computing. R Core Team. R: A Language and Environment for Statistical Computing. Preprint at (2022).

64. Kassambara, A. ggpubr: ‘ggplot2’ Based Publication Ready Plots. Preprint at (2023).

65. Yan, L. ggvenn: Draw Venn Diagram by ‘ggplot2’. CRAN: Contributed Packages Preprint at 10.32614/CRAN.package.ggvenn (2021).

66. Ahlmann-Eltze, C. ggupset: Combination Matrix Axis for ‘ggplot2’ to Create ‘UpSet’ Plots. CRAN: Contributed Packages Preprint at 10.32614/CRAN.package.ggupset (2019).

67. Wagih, O. ggseqlogo: a versatile R package for drawing sequence logos. Bioinformatics 33, 3645–3647 (2017).

68. OpenAI. ChatGPT. Preprint at https://openai.com.

69. Vizcaíno, J. A. et al. ProteomeXchange provides globally coordinated proteomics data submission and dissemination. Nat Biotechnol 32, 223–226 (2014).

