## Supplementary Information for "MAETi: Mild acid elution in a tip enables immunopeptidome profiling from 1E4 cells"

**Document Description:**

The following document contains additional Figures as referenced in the manuscript "MAETi: Mild acid elution in a tip enables immunopeptidome profiling from 1E4 cells".

#### 1. Overview

### 2. Supplementary Figures

The subsequent section contains the supplementary figures (Fig. S#) referenced in the results section of the manuscript.

#### 2.1. Fig. S1: A minimal MAE-based MHC1p enrichment workflow enables immunopeptidomics profiling from 5E4 cells

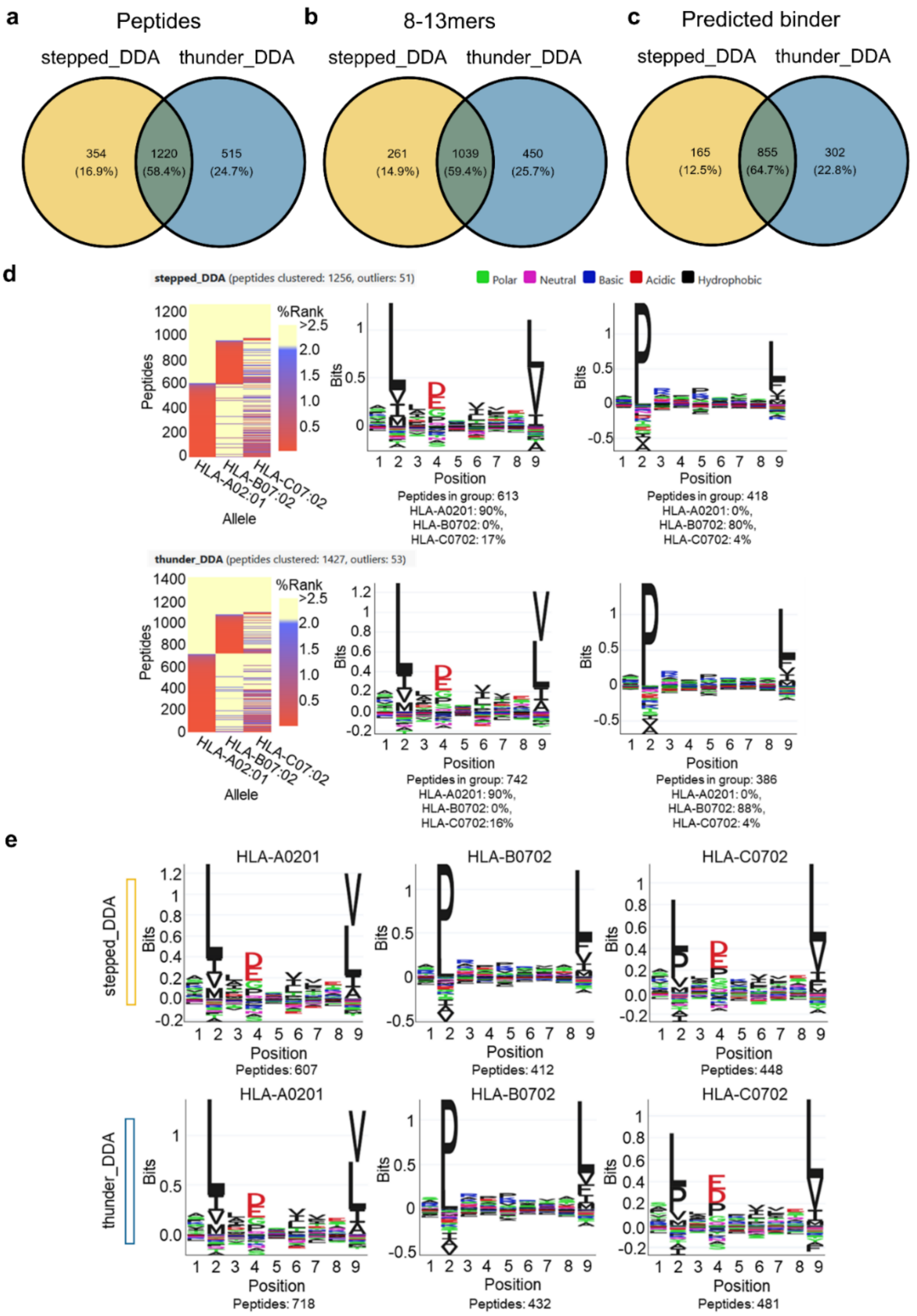

**Fig. S1: A minimal MAE-based MHC1p enrichment workflow enables immunopeptidomics profiling from 5E4 cells (a-e).**
**a-c** Venn diagrams showing the overlap of peptides (**a**), 8-13mers (**b**), and predicted binders (**c**) identified using Stepped-ddaPASEF (“stepped_DDA”, yellow circle) and MHC-tailored Thunder-ddaPASEF (“thunder_DDA”, blue circle). **d, e** unsupervised (**d**) and allele-specific (supervised, **e**) GibbsCluster of identified sequence motifs.

#### 2.2. Fig. S2: Optimized β-Alanine-based MAE Buffer provides cleaner samples and enhances immunopeptidome coverage and reproducibility

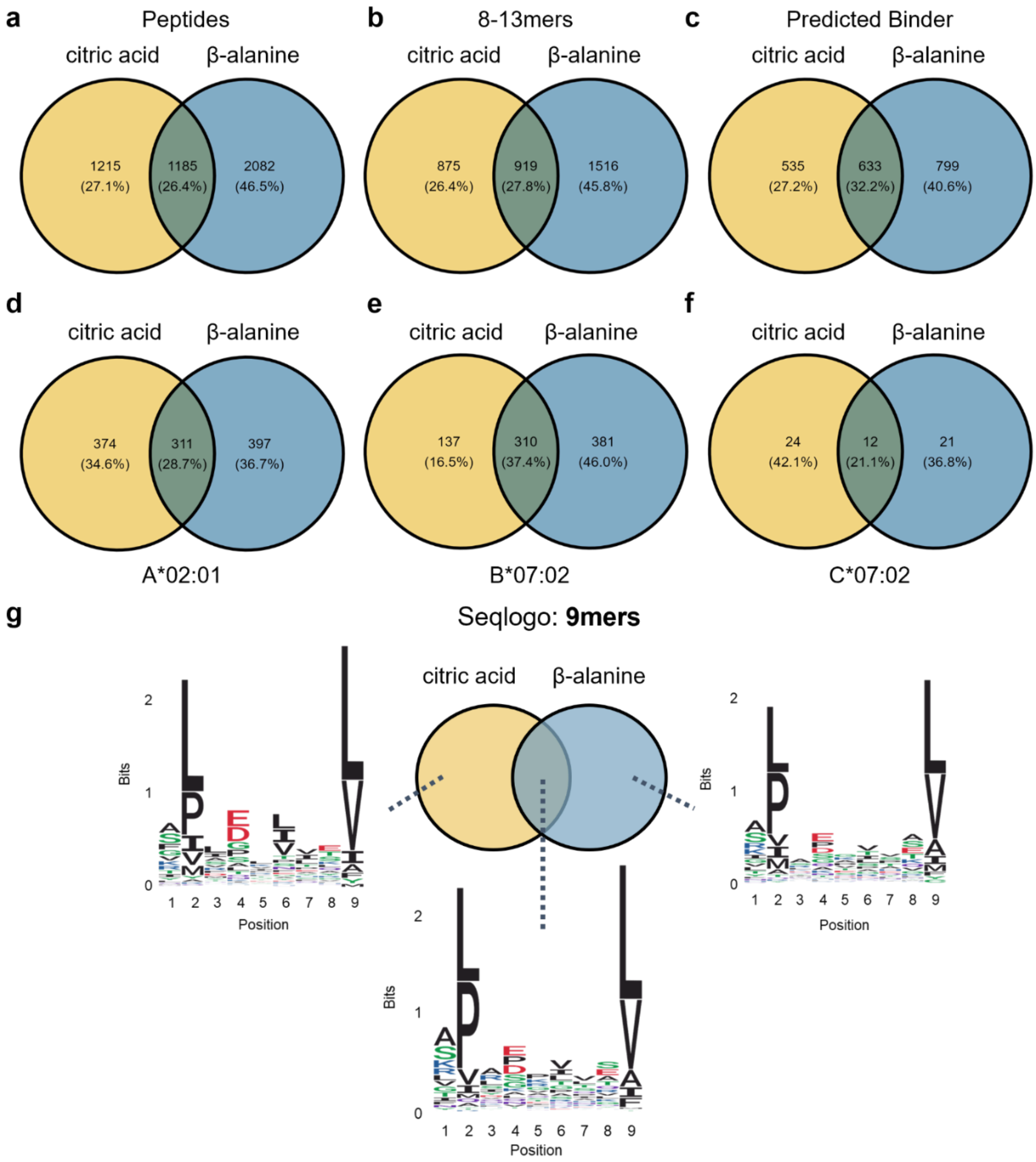

**Fig. S2: 5E4 JY cells per MAETi were prepared using either a Citric acid- or β-Alanine-based MAE buffer (n = 3 each) and analyzed in an Evosep coupled to a timsTOF SCP using Stepped-ddaPASEF.**
**a-f** Venn diagrams showing the overlap of peptides (**a**), 8-13mers (**b**), binders (**c**), and binders by allele (**d-f**) identified using either citric acid or ß-alanine (n = 3, each) as a MAE buffer component. **g** 9mer sequence clustering of predicted binder identified for both citric acid and β-alanine (middle), or exclusively for citric acid (left) or β-alanine (right).

#### 2.3. Fig. S3: β-alanine-based MAE provides a complementary immunopeptidome coverage compared with immunoprecipitation

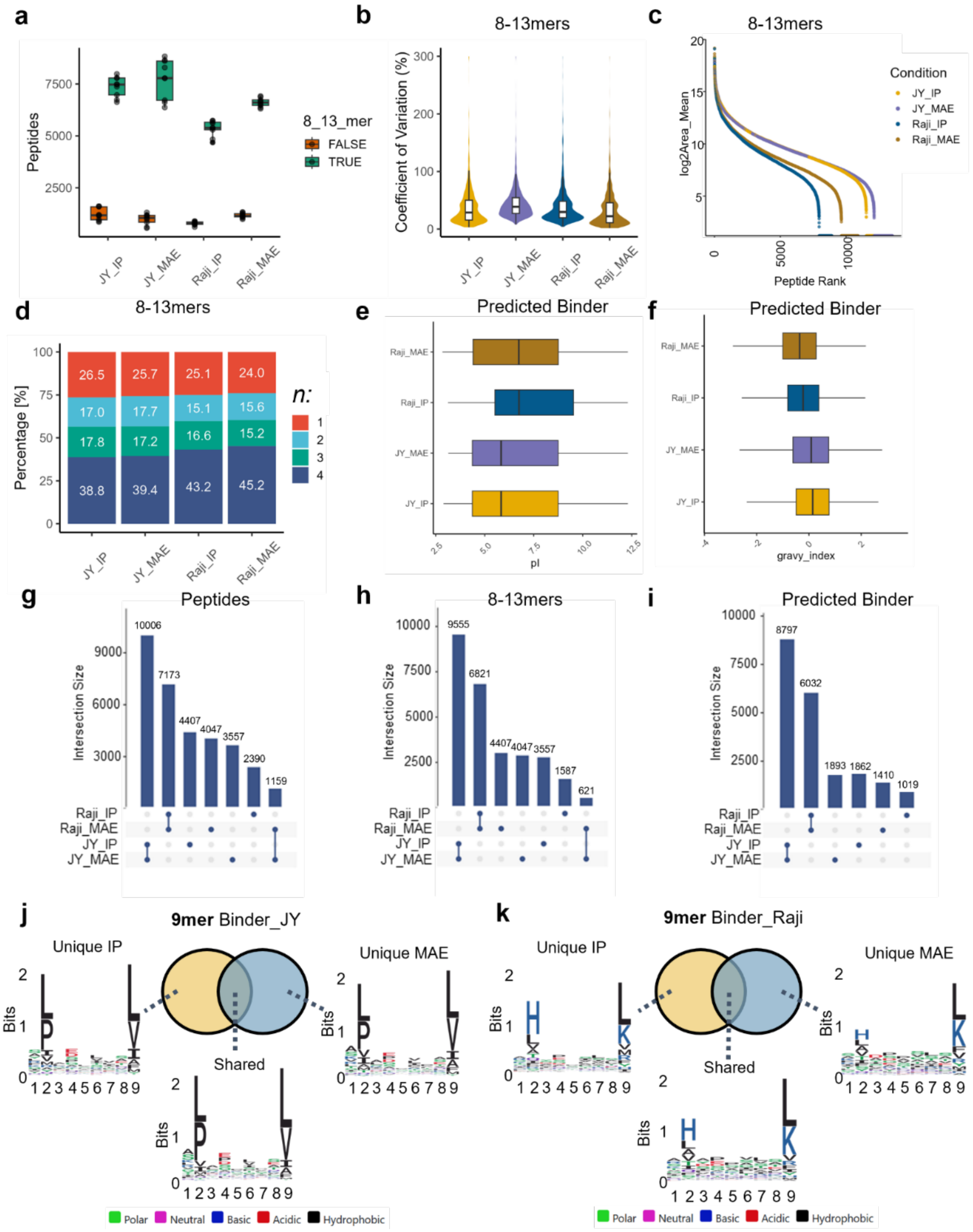

**Fig. S3: β-alanine-based mild acid elution (MAE) provides a complementary immunopeptidome coverage compared with immunoprecipitation (IP) in bulk experiments.**
**a** Boxplot of 8-13mer (“TRUE”) or non-8-13mer identifications (“FALSE”) per cell line and sample preparation technique (center line, median; box limits, upper and lower quartiles; whiskers, 1.5x interquartile range). **b** MS1 Area coefficient of variation (CV) of 8-13mers. **c** Ranked area scatter plot displaying dynamic range of 8-13mer identifications. **d** Data completeness of four out of nine randomly drawn replicates for each condition. **e-f** boxplots displaying the isoelectric point (pI) (**h**) and gravy score (**i**) of predicted binder for each condition. **g-i** Upset plots showing overlaps and uniqueness of peptides (**g**), 8-13mers (**h**), and predicted binder (**i**) between conditions. **j-k** 9mer sequence clustering of predicted binder exclusively identified for either IP (left) or MAE (right) or shared (middle) between both workflows for both JY (**j**) and Raji (**k**).

#### 2.4. Fig. S4: MAETi enables immunopeptidome profiling of SAVs from 1E4 to 1E6 JY cells per sample.

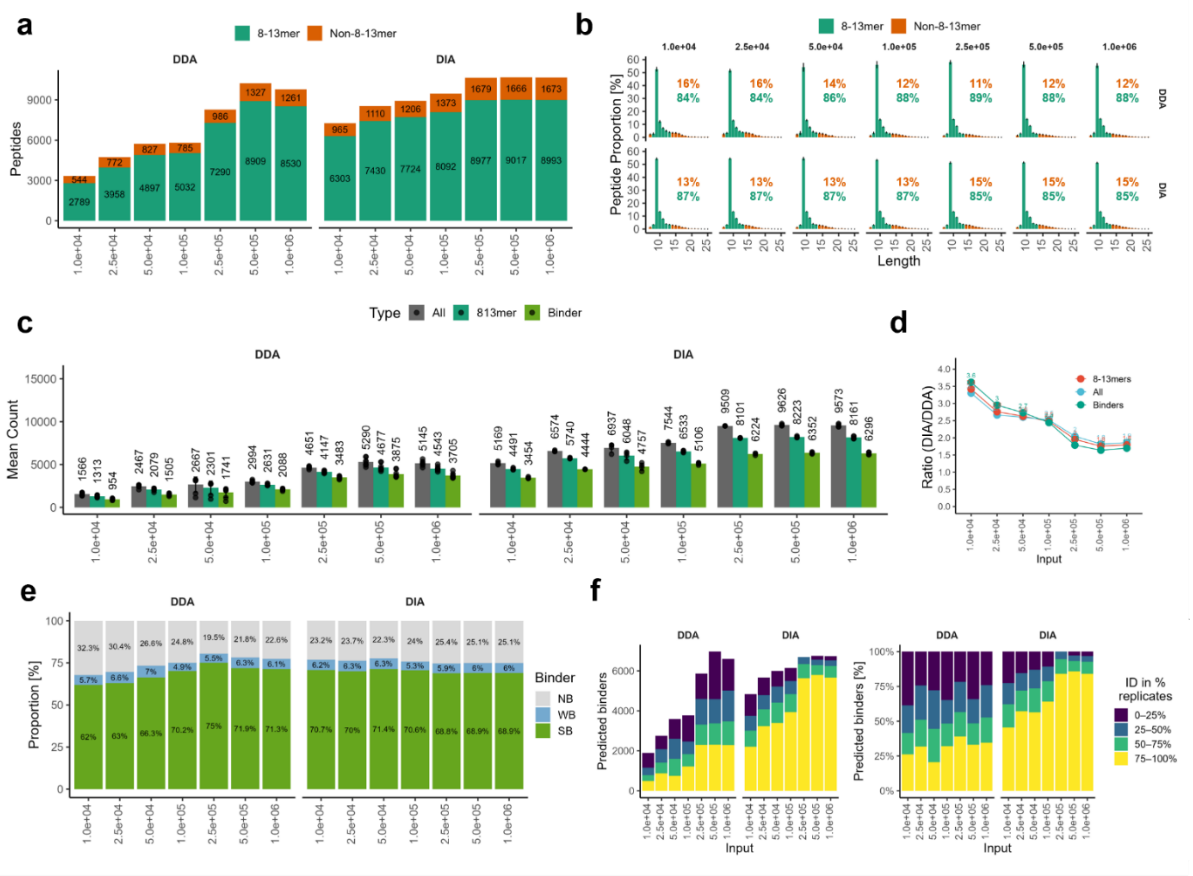

**Fig. S4: MAETi enables immunopeptidome profiling of SAVs from 1E4 to 1E6 JY cells per sample.**
**a** Unique peptides identified in total using Thunder-ddaPASEF ("DDA", n=8) and diaPASEF ("DIA", n=4), divided into 8-13mers (green) and other peptide lengths (orange) across the different input cell numbers. **b** Peptide length distributions with proportion of 8-13mers (green, lower number) and non-8-13mers (orange, upper number) across the different input cell numbers. **c** Peptides, 8-13mers, predicted binder ("Binder") identified on average per sample **d** Ratio of DDA versus DIA identifications for all peptides ("All"), 8-13mers, and predicted binder ("Binder") **e** Proportion of 8-13mers predicted to bind JY MHC alleles (NB, non-binders; WB, weak-binders; SB, strong binders). **f** Data completeness stacked bar chart with total (left) and proportion (right) of identified predicted binders across binned replicates e.g., identified in 0-25% of the replicates for a given input cell number in DDA and DIA, respectively.

#### 2.5. Supplementary Fig. S5-S7: MAETi enables MHC I ligandomics of FACS-sorted immune cells

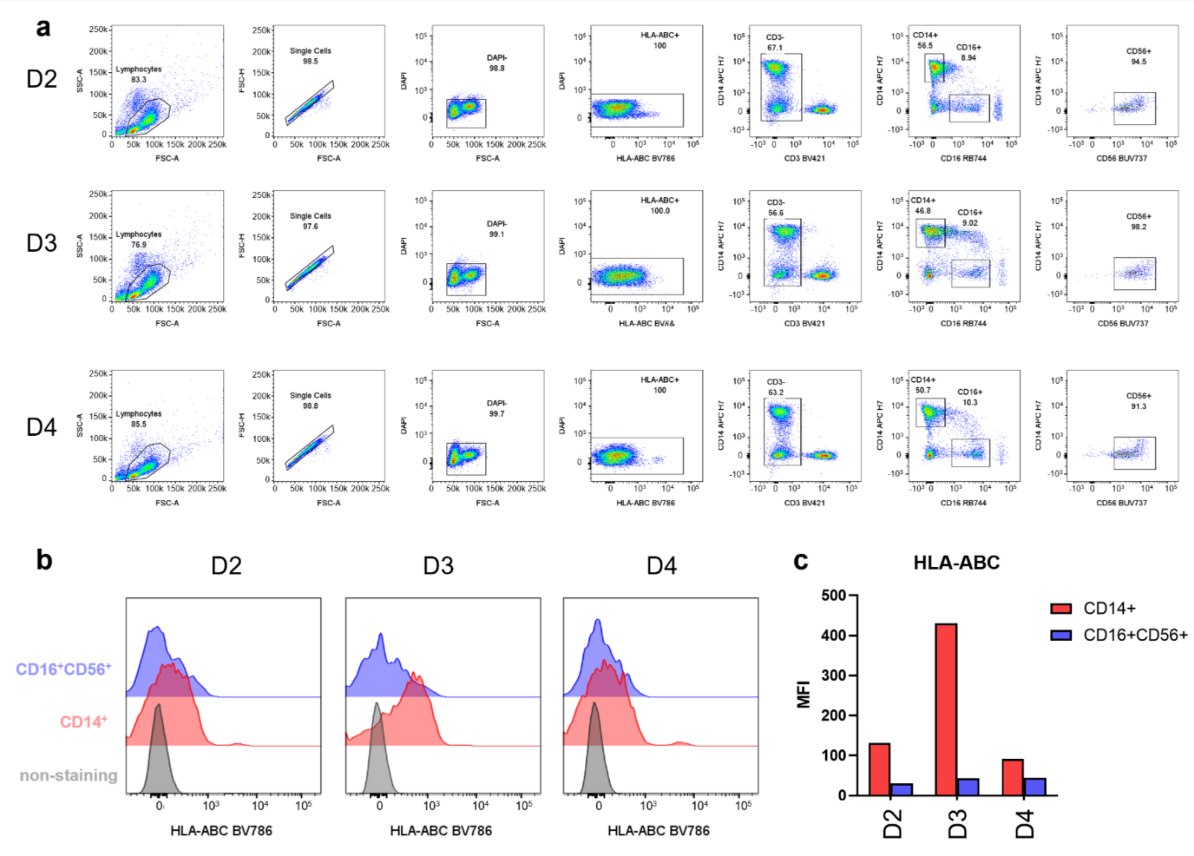

**Fig.S5: MAETi enables immunopeptidome profiling of fluorescence-activated sorted immune cells from human PBMCs**
**a** Exemplary flow cytometric identification of PBMC sub-cell populations for Donor 2 to Donor 4 (D2-D4) stained with DAPI, anti-HLA-ABC, anti-CD3, anti-CD14, anti-CD16 and anti-CD56. **b-c** HLA-ABC expression level versus non-staining in $CD14^+$ and $CD16^+CD56^+$ cells by FACS (b) and bar graph showing mean fluorescence intensity (MFI) of HLA-ABC expression of each donor (c).

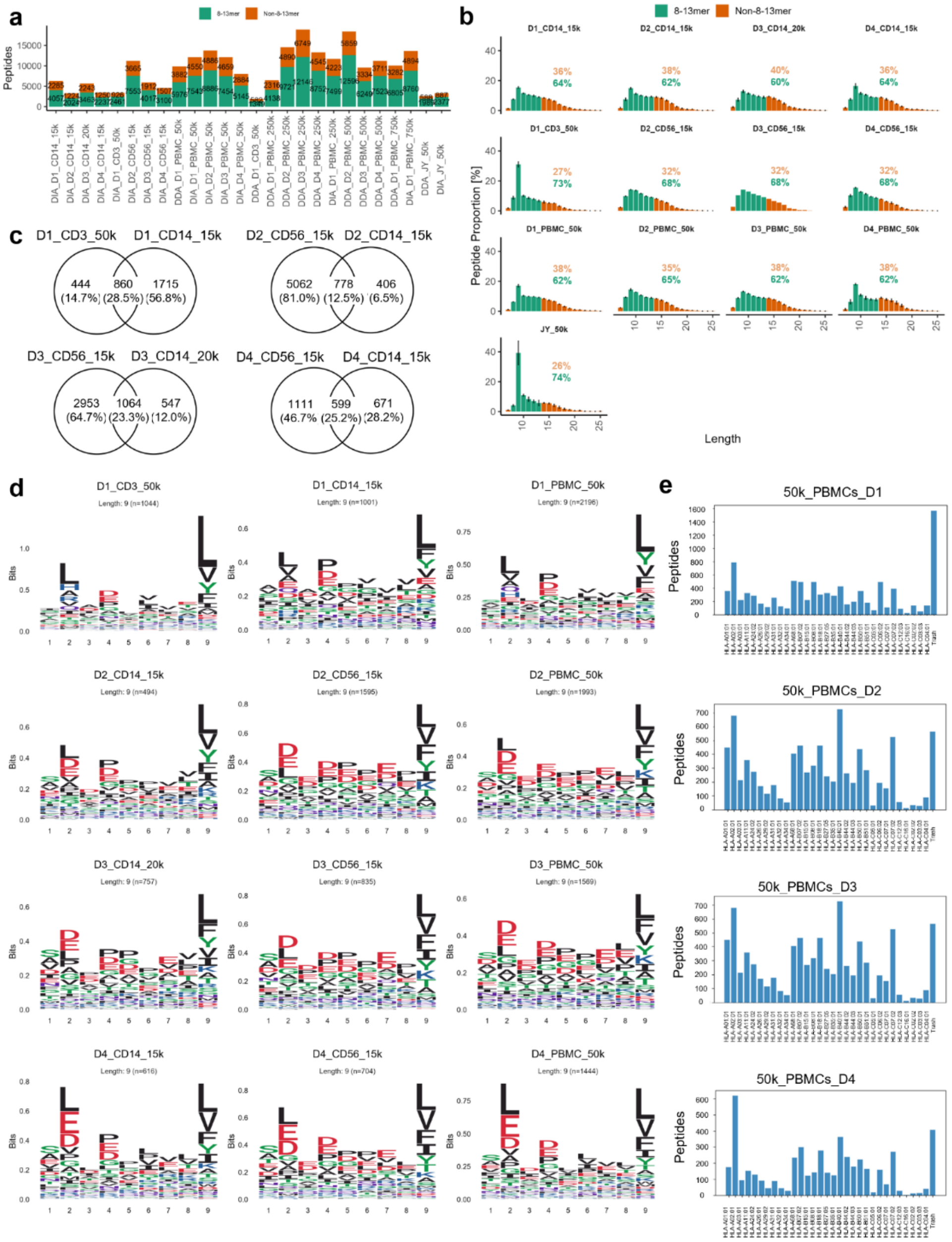

**Fig.S6: MAETi enables immunopeptidome profiling of fluorescence-activated sorted immune cells from human PBMCs**

**a** Unique peptides identified in total using Thunder-ddaPASEF ("DDA") for spectral library generation using 25E4-75E4 PBMCs or diaPASEF on 1.5E4-5E4 isolated cell types and 5E4 whole PBMCs divided into 8-13mers (green) and other peptide lengths (orange). Note that Donor 1 (D1) was 2-digit HLA-typed and, as a control of the MAETi workflow, 5E4 JY cells were processed in parallel and measured both in DDA and DIA **b** Peptide length distributions with proportion of 8-13mers (green, lower number) and non-8-13mers (orange, upper number) for diaPASEF acquisitions. **c** Venn diagrams showing the overlap of 8-13mers detected in at least 60% of the replicates of one condition (i.e., cell type). **d** Unsupervised sequence clustering of the 9mers identified for isolated cell types and PBMCs featuring strong anchor amino acids at position P2 and P9 as well as P4. **e** Bar graphs indicating likely MHC supertypes present in 5E4 whole PBMCs of D1-4 obtained via MHCMotifDecon (v1.0).

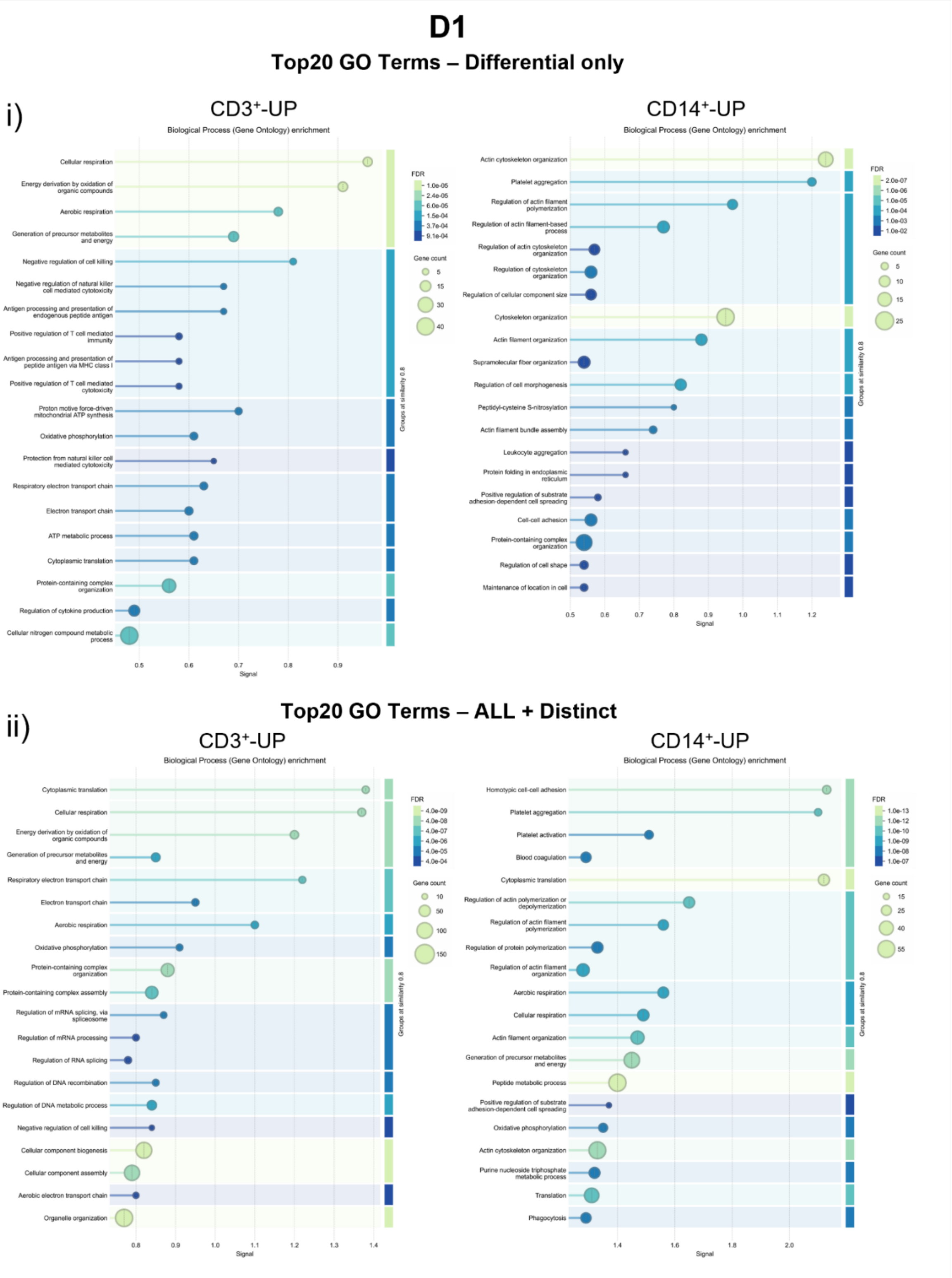

**Fig.S7a: Top 20 Gene Ontology (GO) Terms for D1: MAETi enables immunopeptidome profiling of fluorescence-activated sorted immune cells from human PBMCs**
**i-ii)** Biological processes represented by differentially upregulated 8-13mers (I, upper panel) and differentially upregulated plus distinct 8-13mers (ii, lower panel) identified in at least 60% of the replicates of one condition (i.e., cell type) for CD3$^+$- (left) and CD14$^+$- (right) cells by the strongest signal.

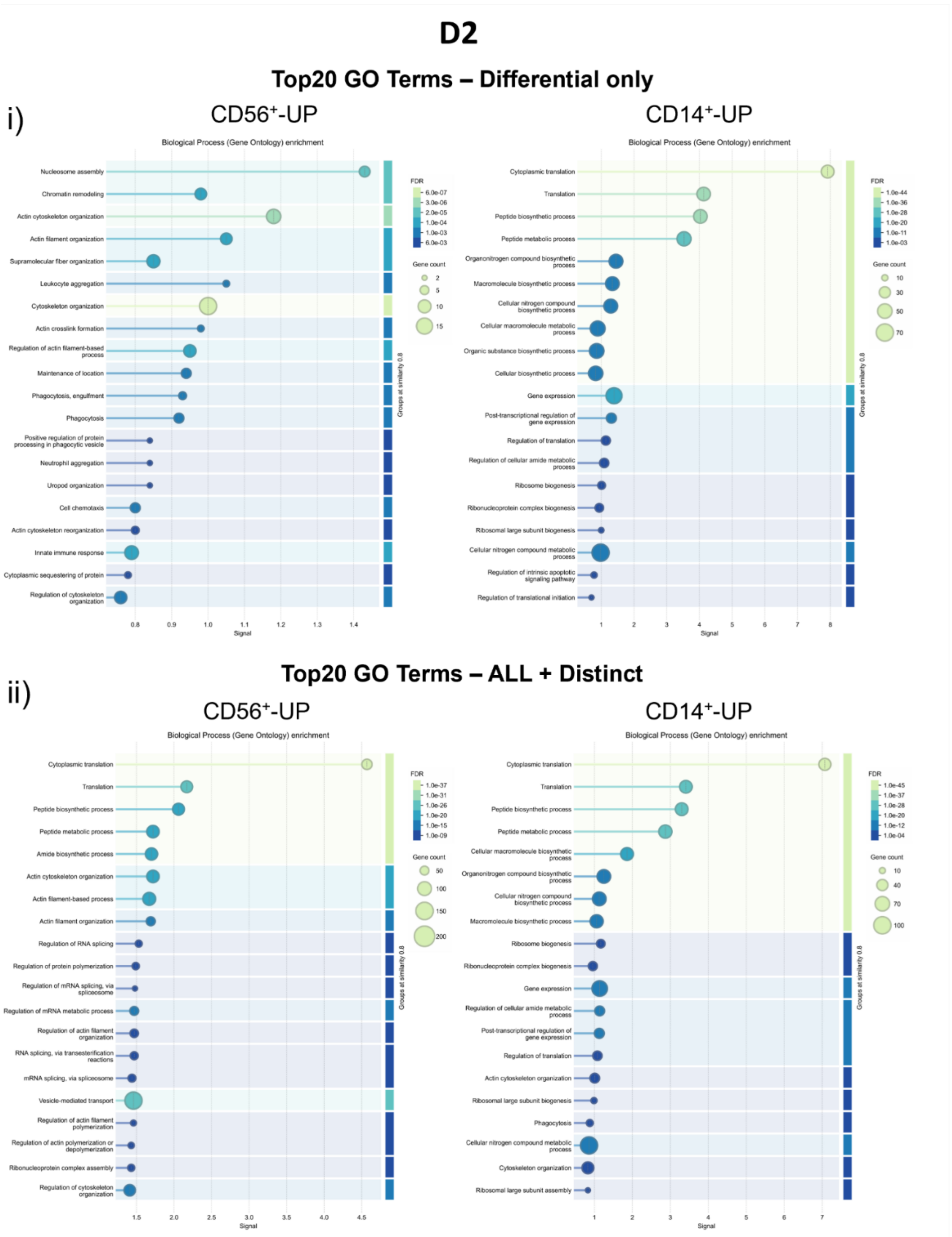

**Fig.S7b: Top 20 Gene Ontology (GO) Terms for D2: MAETi enables immunopeptidome profiling of fluorescence-activated sorted immune cells from human PBMCs**
**i-ii)** Biological processes represented by differentially upregulated 8-13mers (I, upper panel) and differentially upregulated plus distinct 8-13mers (ii, lower panel) identified in at least 60% of the replicates of one condition (i.e., cell type) for $CD16^{+}CD56^{+}$- (left) and $CD14^{+}$- (right) cells by strongest signal.

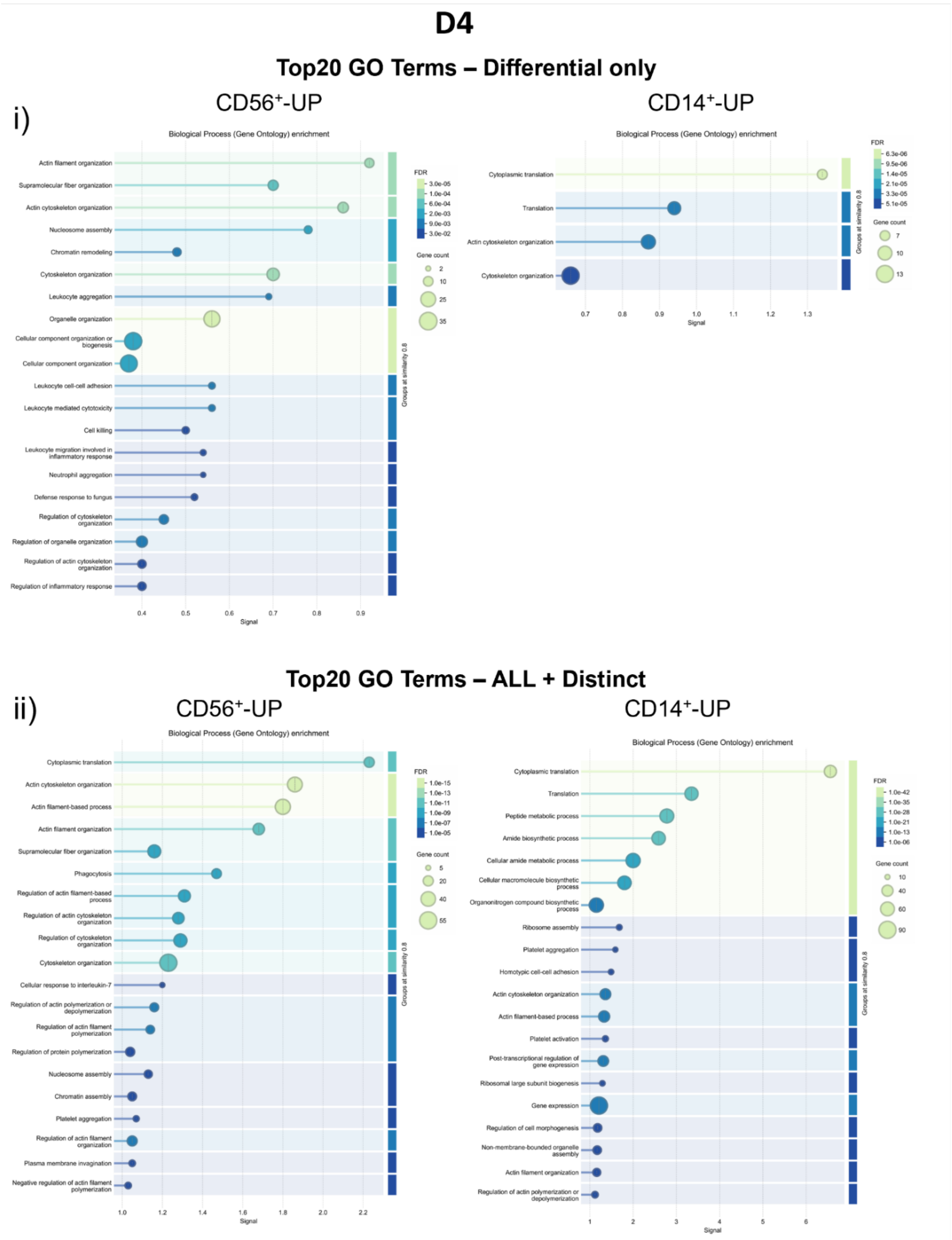

**Fig.S7c: Top 20 Gene Ontology (GO) Terms for D4: MAETi enables immunopeptidome profiling of fluorescence-activated sorted immune cells from human PBMCs**
**i-ii)** Biological processes represented by differentially upregulated 8-13mers (I, upper panel) and differentially upregulated plus distinct 8-13mers (ii, lower panel) identified in at least 60% of the replicates of one condition (i.e., cell type) for CD16⁺CD56⁺- (left) and CD14⁺- (right) cells by the strongest signal.

**Supplementary Data 1. Detailed Sample Overview, MS and Processing Settings**

Excel file with detailed sample overview, MS acquisition parameters, LC-gradient and software processing settings used to generate all figures in this manuscript containing the following worksheets:

- SampleOverview: For each figure comprises a detailed listing of raw file names, LC and MS parameters, processing software, name of uploaded result file, FASTA File used for processing and a reference to settings used during processing (see sheet tabs “PEAKS_Setting” and “FragPipe_Setting”).
- LC_47min: 47 min LC gradient used for the bulk comparison on the nanoElute 2
- PEAKS_Setting: Parameters used for peptide identification in PEAKS Xpro
- FragPipe_Setting: Parameters used for peptide identification in FragPipe v22 and v23.
